# The stability landscape of *de novo* TIM barrels explored by a modular design approach

**DOI:** 10.1101/2020.09.29.319103

**Authors:** Sergio Romero-Romero, Miguel Costas, Daniel-Adriano Silva Manzano, Sina Kordes, Erendira Rojas-Ortega, Cinthya Tapia, Yasel Guerra, Sooruban Shanmugaratnam, Adela Rodríguez-Romero, David Baker, Birte Höcker, D. Alejandro Fernández-Velasco

## Abstract

The ability to design stable proteins with custom-made functions is a major goal in biochemistry with practical relevance for our environment and society. High conformational stability lowers protein sensitivity to mutations and changes in the environment; thus, understanding and manipulating protein stability will expand the applications of *de novo* proteins. Since the (**β/α**)_8_-barrel or TIM-barrel fold is one of the most common functional scaffolds, in this work we designed a collection of stable *de novo* TIM barrels (DeNovoTIMs), using a computational fixed-backbone and modular approach based on improved hydrophobic packing of sTIM11, the first validated *de novo* TIM barrel. DeNovoTIMs navigate a region of the stability landscape previously uncharted by natural TIM barrels, with variations spanning 60 degrees in melting temperature and 22 kcal per mol in conformational stability throughout the designs. Significant non-additive or epistatic effects were observed when stabilizing mutations from different regions of the barrel were combined. The molecular basis of epistasis in DeNovoTIMs appears to be related to the extension of the hydrophobic cores. This study is an important step towards the fine-tuned modulation of protein stability by design.

**Significance Statement:** *De novo* protein design expands our knowledge about protein structure and stability. The TIM barrel is a highly relevant fold used in nature to host a rich variety of catalytic functions. Here, we follow a modular approach to design and characterize a collection of *de novo* TIM barrels and subjected them to a thorough folding analysis. Non-additive effects modulate the increase in stability when different regions of the barrel are mutated, showing a wide variety of thermodynamic properties that allow them to navigate an unexplored region of the stability landscape found in natural TIM barrels. The design of stable proteins increases the applicability of *de novo* proteins and provides crucial information on the molecular determinants that modulate structure and stability.

**One Sentence Summary:** A family of designed TIM barrels with diverse thermodynamic properties shows epistatic effects on its stability landscape.

## Introduction

Proteins are essential macromolecules capable of performing diverse and exquisite biological functions such as nutrient uptake, environmental stimuli sensing, immune protection, energy storage, cellular communication, molecule transportation, or enzymatic reactions. To guarantee such activities, the functional states must act under specific environmental conditions in a relevant time scale, that is, proteins must be “*stable*”. Protein stability is required to maintain functional structures and it enhances the ability of proteins to evolve new properties (*1, 2*). The central role of proteins in the chemistry of life, as well as their increasing application in basic and applied research, implies that the understanding and manipulation of protein stability are both practically and academically relevant.

There are two main indicators of protein conformational stability at equilibrium. One is the difference of free energy between the native and unfolded states at a given temperature (**Δ***G*), which is often obtained by chemical unfolding experiments carried out at 25 °C. In addition, stability is also assessed in the context of thermal unfolding, where the unfolding temperature (*T*_*m*_), the temperature at the midpoint of the transition from native to the unfolded state, is the most common parameter employed to quantify stability. Both the **Δ***G* and *T*_*m*_ parameters, usually determined as criteria for a “*stable*” protein, are related with the enthalpy (**Δ***H*) and heat capacity (**Δ***C*_*P*_) changes through the Gibbs-Helmholtz equation, which describes the variation of **Δ***G* with temperature, the so-called “*stability curve*” of proteins (*3*). Different mechanisms have been proposed to modify the stability curve of proteins (*4*), and numerous studies on natural proteins and their site-directed mutants have been used to rationalize the stability of thermophilic proteins and moreover to engineer thermostability (*5*).

Historically, the design of stable proteins has been one of the main objectives of computational protein design (*6*). Several strategies, such as increasing the hydrophobic area in internal cores, improvement of water-protein interactions, the introduction of disulfide bridges as well as the addition of salt bridges, have been proposed (*7-18*). The design of *de novo* proteins can further enhance our understanding of the physicochemical properties that modulate stability. For example, although folding behavior has been only addressed for very few cases, the kinetic analysis of the folding mechanism of two *de novo* **βα** proteins has revealed complex free energy surfaces (*19, 20*). The fine-tuning of conformational stability, that is, the manipulation of the protein stability curve, is an open challenge for protein design and engineering. Such a goal requires a comprehensive characterization of *de novo* proteins, describing the combination of thermodynamic parameters that can be reached in a particular fold.

Within the different topologies that a protein can adopt, the TIM-barrel or (**β/α**)_8_-barrel fold is one of the most abundant superfolds in nature (*21*). Furthermore, proteomic analysis shows that the TIM-barrel domain is within the mean size of the proteins present in *Escherichia coli* (*22*). Besides, the TIM-barrel fold is one of the most successful topologies used in nature to host catalytic activities. Due to its large variety of functions and its ubiquity in different types of enzymes, the TIM barrel represents a very suitable scaffold for protein function design and engineering (*23*). For these reasons, its construction has been an important objective for protein design over the years (*24-28*). Recently, the successful design of a *de novo* four-fold symmetric TIM barrel was described: the sTIM11 protein (*29*). Considering that sTIM11 presents a sequence distant from the naturally occurring TIM-barrel superfamilies, the potential of the TIM-barrel fold to bear functions is more significant than we know so far. sTIM11 shows a high melting temperature (*T*_*m*_= 80 °C) but low conformational stability (**Δ***G*_25°C_*= ∼*4 kcal mol^-1^) when compared to natural TIM barrels (*29-32*). Since low conformational stability often results in high sensitivity to mutations and changes in the environment, this can limit the design of novel proteins with new functions (*8*). Thus, fine-tuning the stability of the sTIM11 scaffold is a prerequisite to functionalize and generate tailor-made barrels for applications in biochemistry, biotechnology, and medicine. In this work, a fixed-backbone design with a modular approach was used to generate a collection of *de novo* TIM barrels. Their stability landscape and structural properties were characterized in detail increasing our knowledge on how stability can be fine-tuned by design.

## Results and Discussion

### Modular repacking of the TIM-barrel hydrophobic cores

The *de novo* protein sTIM11 is an idealized four-fold symmetric TIM barrel of 184 residues, which was designed to include two cysteines that, however, did not form the intended disulfide bond (Fig. 1). To avoid reactive free thiols, both residues were reverted to the residues in the original four-fold design (C8Q and C181V), resulting in sTIM11noCys. The base design DeNovoTIM0, which is the starting point for all further constructs in this work, additionally contains the changes W34V and A38G in all symmetry-related quarters. These residues are situated in every second **α**/**β**-loop, and in sTIM11, these tryptophan residues are the most highly solvent exposed. While different strategies have been explored to increase protein stability (*8, 18*), here we focused on hydrophobic repacking. The structural analysis suggested two regions to be amenable to improvements in sTIM11, namely the central and the peripheral hydrophobic cores. The interior of the circular sheet forms the central core, whereas the outer face of the strands and the internal face of the helices constitute the peripheral core. In this latter, we identified two regions with internal cavities that are located in the lower and upper parts of the barrel, respectively (Fig. 1). The residues lining the three aforementioned regions were subjected to fixed-backbone Rosetta design according to the flow diagram depicted in Fig. S1.

**Fig. 1.**
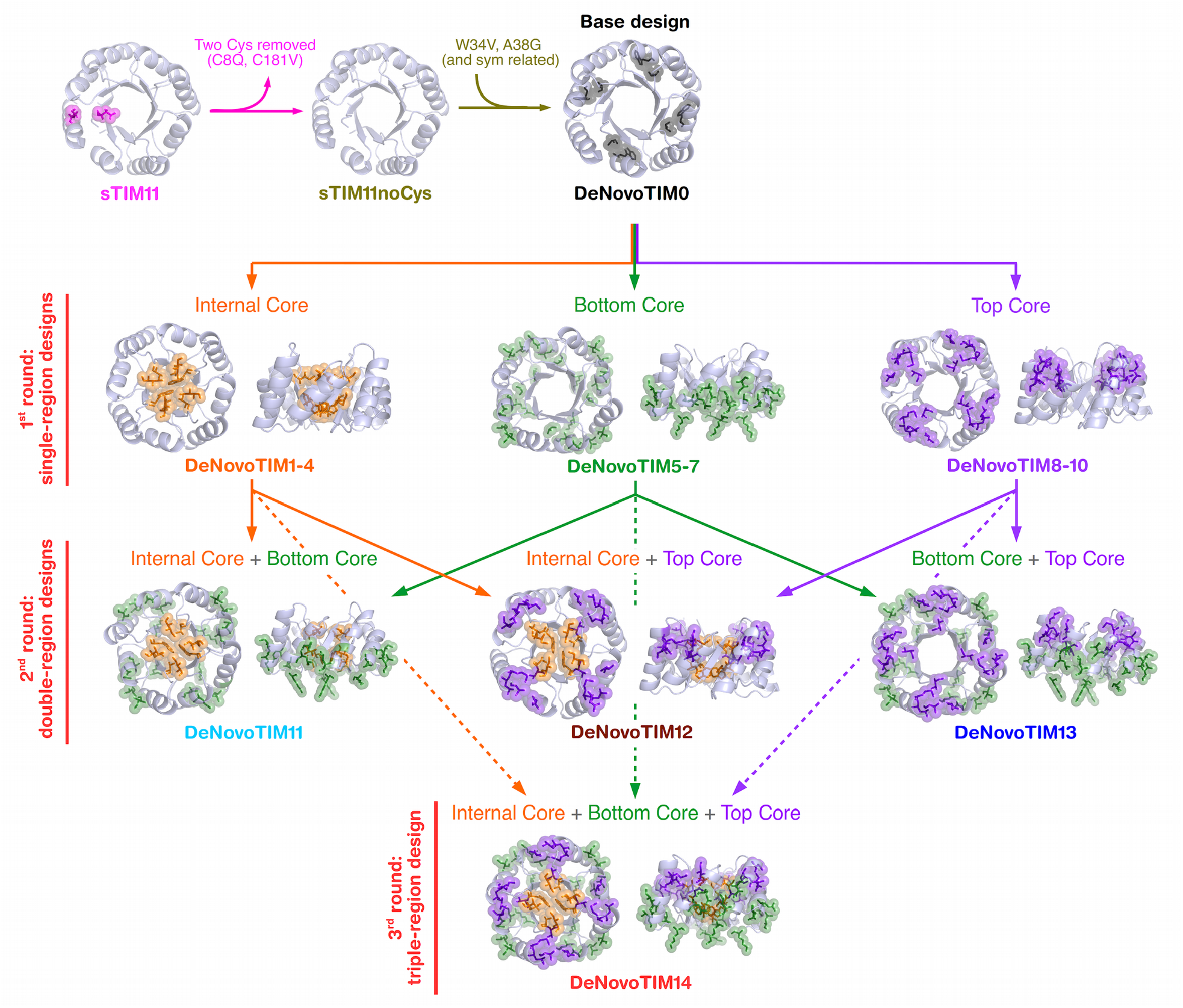
Modular design approach to obtain the DeNovoTIM collection. Cartoon representation of the regions and the corresponding residues modified in each design round. The two cysteine residues present in sTIM11 that were reverted to the corresponding symmetry-related residues in sTIM11noCys are shown in magenta (C8Q and C181V). Mutations W34V and A38G (as well as their 4-fold-symmetry related residues) introduced in DeNovoTIM0 are shown in black. The inner core, formed by the **β**-barrel residues A21, R23, I40, I42, A67, R69, I86, and I88 (as well as their 2 fold-symmetry related residues) is shown in orange. The peripheral bottom core, formed by the N-terminal region of even **β**-strands and the C-region of the flanking **α**-helices, that is, residues Q11, E15, T18, K31, and V34 (as well as their 4-fold-symmetry related residues) is colored green. Peripheral top core situated at the C-terminal region of the odd **β**-strands and the N-terminal region of the flanking **α**-helices formed by residues K2, A5, W6, Y22, S24, and D29 (as well as their 4-fold-symmetry related residues) is shown in purple. All the sequences analyzed in this work are reported in Fig. S2 and tables S1-S2.

Ten designs were selected for characterization in the first round: four with modifications in the internal core (DeNovoTIM1-4) as well as three designs each for the bottom peripheral or outer core (DeNovoTIM5-7) and the top peripheral core (DeNovoTIM8-10) (Fig. S2). For the inner core, no improved designs could be identified when four-fold symmetry was preserved. Therefore, in DeNovoTIM1-4, as well as all of its descendants, only a two-fold symmetry was enforced. An exploratory characterization by circular dichroism (CD) and differential scanning calorimetry (DSC) of these proteins (DeNovoTIMs 1-10) as well as DeNovoTIM0 showed that DeNovoTIM1, DeNovoTIM6, and DeNovoTIM8 were the best designs for each region (Fig. S3 and supporting text).

To test for additivity effects on stability and structure, mutations contained in the best design of each group were combined to generate the following double-region designs: DeNovoTIM11 (DeNovoTIM1 + DeNovoTIM6), DeNovoTIM12 (DeNovoTIM1 + DeNovoTIM8), and DeNovoTIM13 (DeNovoTIM6 + DeNovoTIM8). Finally, in the third design round the mutations of all three regions were combined resulting in DeNovoTIM14 (DeNovoTIM1 + DeNovoTIM6 + DeNovoTIM8). All these proteins as well as sTIM11, sTIM11noCys, and DeNovoTIM0 were characterized in detail (Fig. 1). Information on sequences, and mutations in each design are reported in the supporting information (Fig. S2 and tables S1-S4).

### Physicochemical characterization of DeNovoTIMs

sTIM11, sTIM11noCys, and all DeNovoTIM variants presented the characteristic far-UV CD spectra observed for **α**/**β** proteins (Fig. 2A and Fig. S4). The near-UV CD and intrinsic fluorescence (IF) spectra showed that the aromatic residues are buried from the solvent and structured in the folded state (Fig. 2B-2C and Fig. S5-S6; see supporting text for details). All DeNovoTIMs adopt a monomeric and compact shape as revealed by the invariant value of the Stokes radius determined by analytical size exclusion chromatography over a twenty-fold protein concentration range (table S5). DeNovoTIM0 is also monomeric but shows a Stokes radius (26.1 ± 0.3 Å) slightly higher than that expected for a compact protein of this size, but still far away from the expected value for an unfolded conformation (22.5 ± 1.0 Å and 42.0 ± 1.0 Å, respectively; *33*). This is in agreement with the red shift in the IF spectra and suggests a slightly expanded conformation for DeNovoTIM0.

**Fig. 2.**
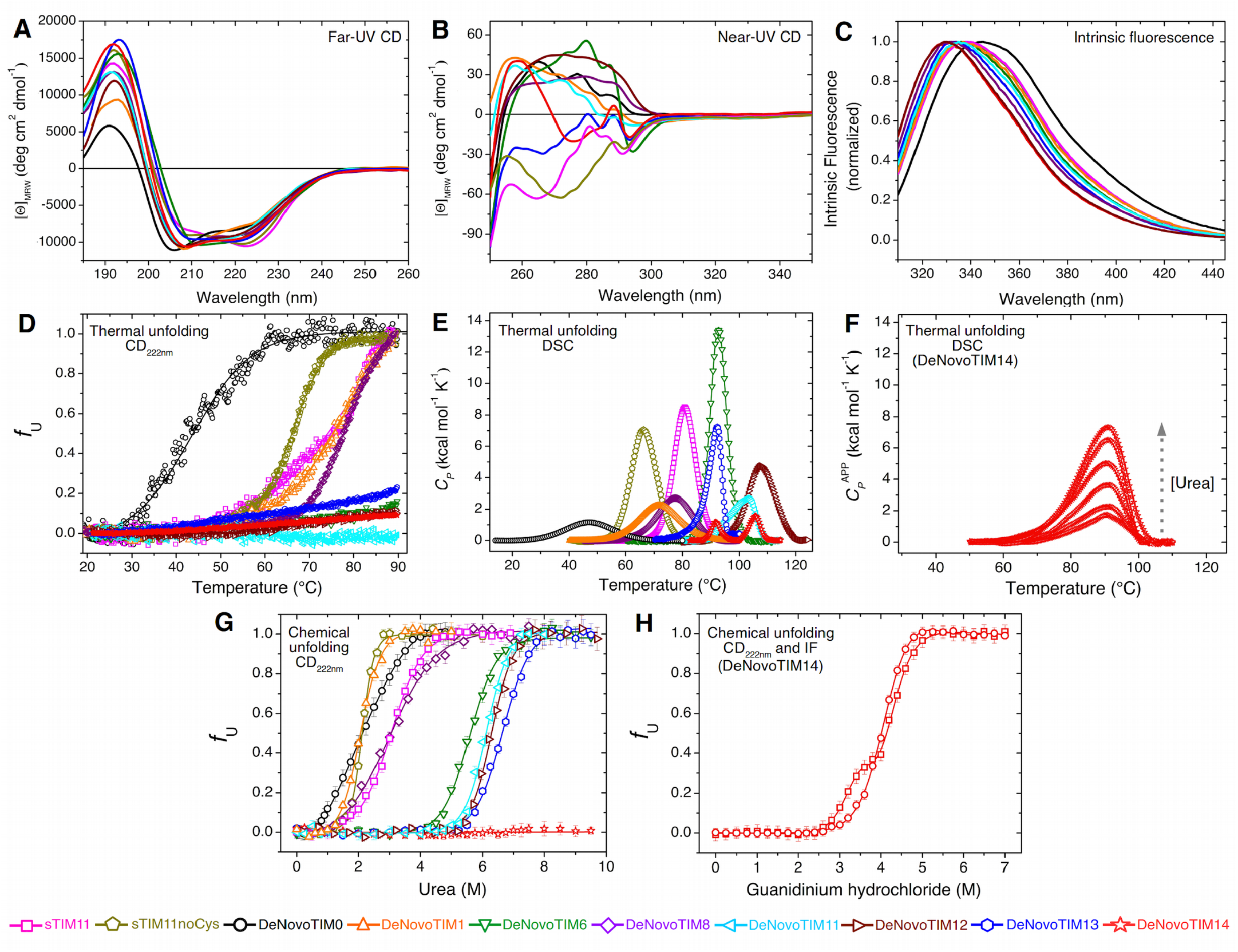
Conformational properties and equilibrium unfolding of DeNovoTIMs. **A**) Far-UV CD spectra. **B**) Near-UV CD spectra. **C**) Intrinsic Fluorescence (IF) spectra (λ_exc_= 295 nm). **D**) Thermal unfolding followed by CD_222nm_ (scan rate: 1.5 K hr^-1^). **E**) Differential Scanning Calorimetry (DSC) endotherms (scan rate: 1.5 K hr^-1^; for easy comparison, the physical and chemical baselines have been subtracted). **F**) DSC endotherms of DeNovoTIM14 in the presence of increasing concentrations of urea (2.0 to 6.0 M) from bottom to top (scan rate: 1.5 K hr^-1^). For clarity, in panels E and F only a small part of the pre- and post-transition baselines are shown. **G**) Chemical unfolding using urea and followed by CD (notice that DeNovoTIM14 does not unfold with urea). **H**) Chemical unfolding induced by guanidinium hydrochloride for DeNovoTIM14 (squares: CD, circles: IF).

Thermal unfolding was then studied by CD and DSC (Fig. 2D-2E). All DeNovoTIMs showed cooperative transitions with a remarkably broad range of *T*_*m*_ values, from 47 °C (DeNovoTIM0) to 109 °C (DeNovoTIM12) (table 1); indeed at 90 °C many of the proteins still showed secondary and tertiary structure (Fig. S4B and Fig. S5B). All DeNovoTIMs, except DeNovoTIM13 and DeNovoTIM14, showed thermal unfolding reversibility (Fig. S7) and were well fitted to the two-state model (N**⇋**U) (Fig. S8 and table 1). This is remarkable because the temperature-induced unfolding of natural proteins of this size, particularly TIM barrels, is usually not reversible (*22, 30*). DeNovoTIM14 showed two endotherms, suggesting the presence of an unfolding intermediate (Fig. S8I), For DeNovoTIM13 and DeNovoTIM14, endotherms were well-fitted to an irreversible two-state mechanism (N**→**F) giving activation energies (*E*_*act*_) of 120.3 and 37.4 kcal mol^-1^ (table 1), respectively, resulting in very different kinetic stabilities (Fig. S9-S10 and supporting text).

**Table 1.** Thermodynamic properties of DeNovoTIMs.

| *de novo*<br>TIM barrel | Type of modular design | Thermal unfolding (by DSC)<sup>a</sup> |  |  |  |  |  | Chemical unfolding (by CD and IF)<sup>e</sup> |  |  |
| --- | --- | --- | --- | --- | --- | --- | --- | --- | --- | --- |
|  |  | *T<sub>m</sub>*<br>(°C) | Δ*H*<br>(kcal mol<sup>-1</sup>) | Δ*H*<sub>85°C</sub><br>(kcal mol<sup>-1</sup>)<sup>b</sup> | Δ*C<sub>p</sub>*<br>(kcal mol<sup>-1</sup> K<sup>-1</sup>) | Δ*H<sub>vH</sub>* / Δ*H* | Global<br>Thermodynamic<br>stability<br>(kcal K mol<sup>-1</sup>)<sup>c</sup> | Δ*G*<sub>25°C</sub><br>(kcal mol<sup>-1</sup>) | *m*<br>(kcal mol<sup>-1</sup> M<sup>-1</sup>) | D<sub>[1/2]</sub><br>(M) |
| sTIM11 | First reported *de novo* TIM barrel | 80.0 ± 0.2 | 92.9 ± 1.3 | 104.0 ± 1.8 | 2.19 ± 0.09 | 0.99 ± 0.05 | 279.6 | 4.8 ± 0.3 | 1.34 ± 0.06 | 3.1 |
| sTIM11noCys | sTIM11 without cysteines | 65.6 ± 0.1 | 82.0 ± 0.6 | 127.8 ± 2.2 | 2.36 ± 0.08 | 0.99 ± 0.03 | 175.8 | 3.2 ± 0.2 | 2.03 ± 0.10 | 1.9 |
| DeNovoTIM0 | Base design | 47.0 ± 0.2 | 24.7 ± 0.6 | 41.4 ± 2.9 | 0.44 ± 0.06 | 1.09 ± 0.08 | 60.5 | 1.5 ± 0.1 | 0.76 ± 0.02 | 2.1 |
| DeNovoTIM1 | Single-region design:<br>Internal core | 71.0 ± 0.4 | 48.1 ± 0.8 | 57.8 ± 1.5 | 0.69 ± 0.05 | 0.98 ± 0.05 | 225.9 | 3.8 ± 0.1 | 1.87 ± 0.10 | 2.0 |
| DeNovoTIM6 | Single-region design:<br>Bottom cavity | 92.3 ± 0.1 | 124.9 ± 1.5 | 107.5 ± 1.1 | 2.38 ± 0.06 | 1.03 ± 0.02 | 542.2 | 7.9 ± 0.2 | 1.51 ± 0.08 | 5.6 |
| DeNovoTIM8 | Single-region design:<br>Top cavity | 77.3 ± 0.3 | 51.9 ± 0.8 | 61.1.81.3 | 1.19 ± 0.07 | 0.95 ± 0.09 | 165.2 | 2.5 ± 0.2 | 0.85 ± 0.09 | 2.9 |
| DeNovoTIM11 | Double-region design:<br>Internal core + bottom cavity<br>(DeNovoTIM1 + DeNovoTIM6) | 103.5 ± 0.2 | 61.7 ± 2.4 | 49.3 ± 0.9 | 0.67 ± 0.14 | 1.02 ± 0.09 | 523.2 | 8.4 ± 0.4 | 1.75 ± 0.08 | 6.0 |
| DeNovoTIM12 | Double-region design:<br>internal core + top cavity<br>(DeNovoTIM1 + DeNovoTIM8) | 108.8 ± 0.3 | 79.3 ± 1.9 | 62.2 ± 1.0 | 0.72 ± 0.08 | 1.01 ± 0.08 | 791.5 | 10.9 ± 0.2 | 1.77 ± 0.03 | 6.2 |
| DeNovoTIM13 | Double-region design:<br>bottom cavity + top cavity<br>(DeNovoTIM6 + DeNovoTIM8) | 92.8 ± 0.4 | 46.7 ± 4.5 | ND<br>*E<sub>act</sub>*: 120.3 ± 2.8 kcal mol<sup>-1</sup> <sup>d</sup> |  |  | ND | 9.5 ± 0.2 | 1.54 ± 0.03 | 6.6 |
| DeNovoTIM14 | Triple-region design:<br>internal core + bottom cavity + top cavity<br>(DeNovoTIM1 + DeNovoTIM6 + DeNovoTIM8) | 91.5 ± 0.1 | 5.4 ± 0.2 | ND<br>*E<sub>act</sub>*: 37.4 ± 0.5 kcal mol<sup>-1</sup> <sup>d</sup> |  |  | ND | Tot: 23.6 ± 2.0<br>N to I: 12.7 ± 1.9<br>I to U: 10.9 ± 0.6 | N to I: 4.3 ± 0.9<br>I to U: 2.5 ± 0.1 | N to I: 3.2<br>I to U: 4.4 |
|  |  | 105.3 ± 0.1 | 8.6 ± 0.4 |  |  |  |  |  |  |  |

<sup>a</sup> The thermodynamic parameters reported are the average of ten experiments carried out at different protein concentrations (0.25-2.5 mg mL<sup>-1</sup>; Fig. S8 and Fig. S11), ± indicate the standard deviation calculated from these 10 experiments.
<sup>b</sup> $\Delta H$ at 85 °C ( $\Delta H_{85^\circ\text{C}}$ ) was calculated using the experimental $\Delta H$ and $\Delta C_p$ values as indicated in the experimental section.
<sup>c</sup> The global thermodynamic stability was calculated from the area of the stability curve evaluated between 0 °C and $T_m$ (Fig. 3B).
ND: Not determined due to irreversibility in the thermal unfolding. Instead, activation energy ( $E_{\text{act}}$ ) was calculated from an irreversible two-state mechanism.
<sup>d</sup> $E_{\text{act}}$ value and ± are the average and standard deviation, respectively, from three different calculation methods (Fig. S9 and Fig. S10).
<sup>e</sup> ± indicate the standard error from global fitting (Fig. S13).

For DeNovoTIMs with a reversible thermal unfolding, the observed unfolding **Δ***H* and **Δ***C*_*P*_ also vary greatly (table 1); for some DeNovoTIMs these values are similar to the ones expected for a protein of 184 residues, whereas for others they are smaller (**Δ***H*= 128.4 ± 3.5 kcal mol^-1^ and **Δ***C*_*P*_= 2.6 ± 0.04 kcal mol^-1^ K^-1^, according to parametric equations reported in *34*). The **Δ***H* values observed for the first and second design rounds (0.24 to 0.64 kcal mol^-1^ residue^-1^) are similar to those reported for natural monomeric TIM barrels (0.25 to 0.67 kcal mol^-1^ residue^-1^).

In order to obtain reliable **Δ***C*_*P*_ values, multiple DSC experiments were performed at ten different protein concentrations (Fig. S8 and table 1), obtaining **Δ***C*_*P*_ values which were independent of protein concentration and varied in a small standard deviation range (Fig. S11). A decrease in **Δ***C*_*P*_ has been shown to result from residual structure in the unfolded state (*35*). This is observed in the far-UV CD spectra of those DeNovoTIMs that are unfolded at 90 °C. In addition, the low **Δ***H* of DeNovoTIM14 increases in the presence of urea (Fig. 2F and Fig. S10). These results suggest that for some DeNovoTIMs, the reason for the low **Δ***H* and **Δ***C*_*P*_ is likely the high content of residual structure in the unfolded state (supporting text).

Stability at 25 °C was studied by chemical unfolding with urea or GdnHCl. Except for DeNovoTIM14, all designs were completely unfolded in 9.0 M urea (Fig. S4-S6). Unfolding and refolding transitions are coincident and the signal does not change after incubation for 12 hours, i.e. chemical unfolding is reversible and in equilibrium under the experimental conditions. For all DeNovoTIMs, except for DeNovoTIM14, CD and IF curves were monophasic, cooperative, coincident, and well globally-fitted to a two-state N**⇋**U model, indicating the absence of populated intermediates (Fig. 2G and Fig. S12-S13). DeNovoTIM14 presented a different behavior; no changes in CD or IF signal were observed in the presence of urea (Fig. 2G), even after incubation for 5 days. CD and IF spectra indicate that at 9.0 M urea DeNovoTIM14 presents native-like properties (Fig. S4-S6).

When chemical unfolding was carried out with GdnHCl, unfolding transitions were reversible and at equilibrium with this denaturant. IF data showed a monophasic transition in the 3-5 M GdnHCl range, while CD detected the presence of an unfolding intermediate between 3-4 M GdnHCl (Fig. 2H). Both traces were globally fitted to a three-state model with an intermediate: N**⇋**I**⇋**U (Fig. S13). All the selected first-and second-round designs presented a **Δ***G* at 25 °C higher than DeNovoTIM0, whereas the triple-design, DeNovoTIM14, showed a pronounced increase in stability (**Δ***G*_Tot_= 23.6 kcal mol^-1^; table 1). For DeNovoTIM14, the stability change related to the loss of the native state (**Δ***G*_N-I_= 12.7 kcal mol^-1^) is higher than the **Δ***G* of second-round designs, whereas the stability of the intermediate is similar to them (**Δ***G*_I-U_= 10.9 kcal mol^-1^). For three-state folders, the change in free energy from the native to the intermediate state (**Δ***G*_N-I_) has been termed “the relevant stability” because the intermediate is expected to be non-functional, whereas the stability change from the intermediate to the unfolded state (**Δ***G*_I-U_) is designated as “residual stability” (*36*).

*m* values (*m*= ∂**Δ***G*/ ∂[denaturant]) are proportional to the surface area exposed to the solvent upon unfolding (**Δ**ASA); likewise, the buried area correlates very strongly with the number of residues (*37*). The *m* value calculated from the sTIM11noCys structure is in excellent agreement with the experimentally determined one (2.15 vs. 2.03 kcal mol^-1^ M^-1^). For all the other DeNovoTIMs, the *m* value is similar to those observed for natural proteins with the same size, except for DeNovoTIM0 and DeNovoTIM8 where *m* decreases significantly, indicating that the native structure may not be completely well-packed or that the unfolded state has residual secondary structure (table 1). Although residual structure in the unfolded state is not clearly observed in CD spectra in 9.0 M urea (Fig. S4-S5), the persistence of native-like structure could be present at high urea concentration and not be identified by the techniques used here, as it has been reported for other proteins (*38, 39*).

The modular design approach used in this work improved both **Δ***G* and *T*_*m*_ substantially and hence produced significantly more stable proteins, particularly in the second- and third-round designs. In this context, it is worth mentioning that over the years the combination of stabilizing mutations has been considered an effective strategy to enhance the stability of small proteins (*36, 40-43*). Previous work on small globular proteins with optimized hydrophobic cores and interactions on the surface exhibited increased thermal stability by up to 30 degrees (*9, 11, 15*). Extending these strategies from point mutants to regions appears to be useful for bigger folds such as the TIM barrel. In what follows, using the thermal and chemical unfolding data described above, the thermodynamic properties underlying the stability of DeNovoTIMs are analyzed.

### Global thermodynamic stability and non-additive effects of DeNovoTIMs

As observed in natural proteins, the *m* values obtained from the chemical unfolding of sTIM11, sTIM11noCys, DeNovoTIM0, DeNovoTIM6, and DeNovoTIM8 correlate with their **Δ***C*_*P*_ values determined by thermal unfolding (Fig. 3A), likely because both depend on the **Δ**ASA upon unfolding. In contrast, **Δ***C*_*P*_ values obtained for DeNovoTIM1, DeNovoTIM11, and DeNovoTIM12 are much lower than those expected from the reported correlation between *m* values and **Δ***C*_*P*_ (Fig. 3A). According to the Rosetta models and the native state structures (see below), these differences are not exclusively due to properties of the native state since the calculated **Δ**ASA is close to the expected value for the size of DeNovoTIMs (17 135 A^2^; *37*). This suggests that the unfolded state reached at high temperatures is more structured than the one obtained by chemical unfolding.

The fact that many DeNovoTIMs show reversible temperature-induced unfolding allowed the assessment of their stability curves using the thermodynamic parameters obtained by DSC data (Fig. 3B). The **Δ***G* values at 25 °C are in excellent agreement with those obtained from chemical unfolding experiments. According to the Gibbs-Helmholtz equation, conformational stability is modulated by changes in *T*_*m*_, **Δ***H*, and **Δ***C*_*P*_. For natural TIM barrels, it has been observed that changes in the stability curve are influenced mainly by modifying one or two of those parameters (*30, 31*). In contrast, the DeNovoTIMs differ in all three parameters. Increasing **Δ***H* is the most commonly found mechanism for stabilization of thermophilic proteins (*5*) and is also the most often exploited mechanism for engineering protein stability (*7, 40*). In DeNovoTIMs, this mechanism is used in all proteins but is especially important in DeNovoTIM6, that has the highest **Δ***H* and, therefore, a **Δ***G* higher than DeNovoTIM0, indicating an enthalpy-driven stabilization (Fig. 3B). Nevertheless, in the absence of a high-resolution structure (see below), it is difficult to rationalize how enthalpic stabilization was achieved in DeNovoTIM6 because considerable structural rearrangements take place when new interactions are introduced or molecular strain is removed. **Δ***C*_*P*_ determines the magnitude of the curvature of the stability curve so that changes in this parameter triggers a more or less flattened curve. A decrease in **Δ***C*_*P*_ has been postulated as a mechanism for thermostabilization (*35, 44*). For DeNovoTIMs, the reduction in **Δ***C*_*P*_ combined with an increase in **Δ***H* is the reason for the increase in both *T*_*m*_ and stability at 25 °C. The results presented here indicate that, as observed for natural proteins, in addition to the native state, the unfolded ensemble plays an important role in shaping the stability curve and should be considered in protein design.

**Fig. 3.**
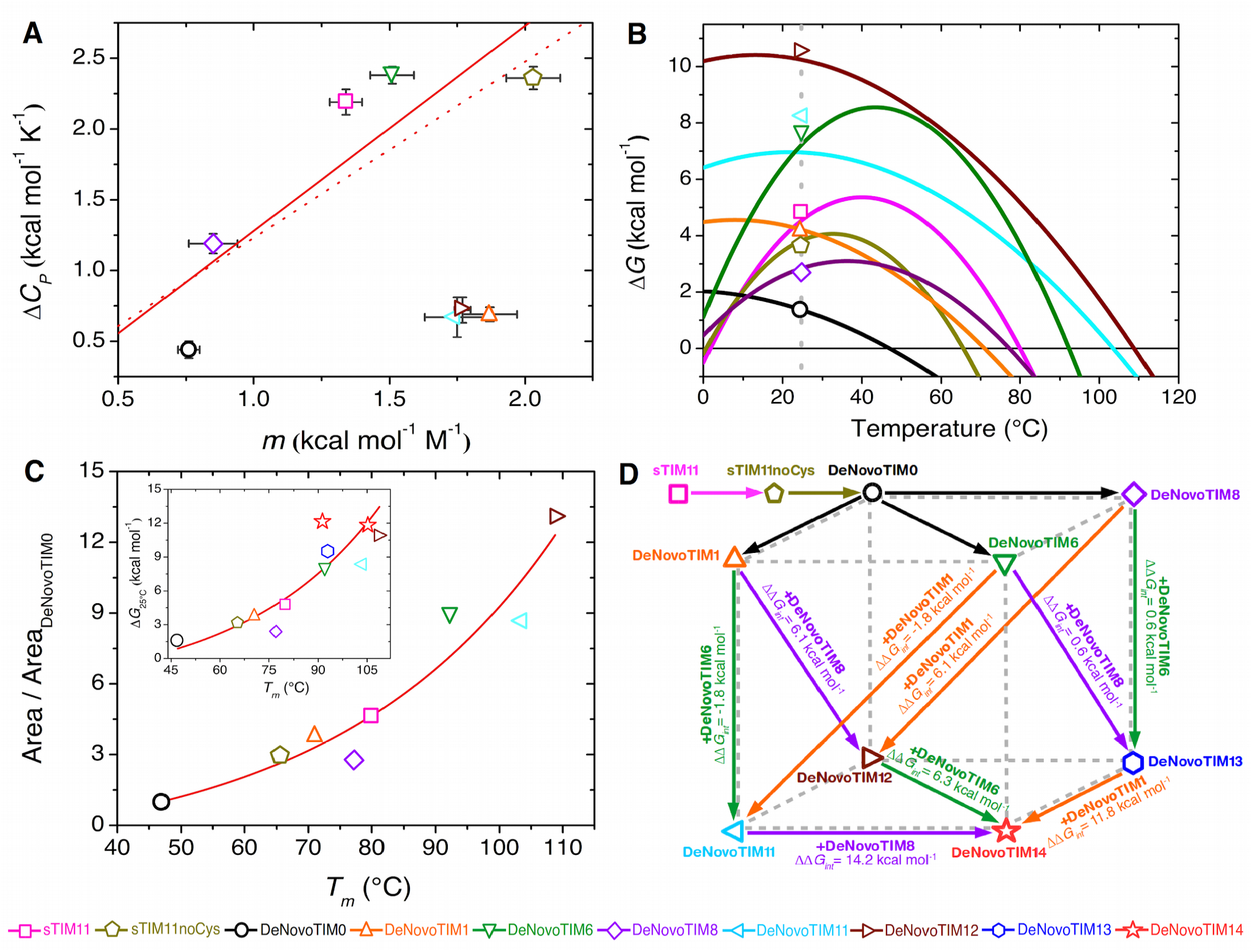
Stability and energetic coupling in DeNovoTIMs. **A**) Correlation between two parameters which are proportional to the exposed surface area: *m* value from chemical unfolding and **Δ***C*_*P*_ from temperature-induced unfolding (solid line: linear regression excluding DeNovoTIM1, DeNovoTIM11, and DeNovoTIM12 data; R^2^: 0.76. Dotted line: correlation reported by *37*). **B**) Stability curves calculated from DSC data (lines) using the Gibbs-Helmholtz equation (open symbols show **Δ***G* values determined by chemical unfolding at 25 °C. Grey dashed line indicates 25 °C). **C**) Correlation between the relative global thermodynamic stability (Area/Area_DeNovoTIM0_) and thermostability (*T*_*m*_) (R^2^: 0.93). Inset: correlation between **Δ***G* at 25 °C determined by chemical unfolding and *T*_*m*_ (R^2^: 0.87). For DeNovoTIM14, where two transitions were found, it was assumed that the one observed at lower [GdnHCl] corresponds to the lower *T*_*m*_. **D**) Thermodynamic cube showing the coupling energy (**ΔΔ***G*_*int*_) between different regions of DeNovoTIMs. **ΔΔ***G*_*int*_ values were calculated from the double-mutant cycles shown in Fig. S14. **ΔΔ***G*_*int*_ values between single-region mutants are depicted as colored arrows from the top face to the bottom face. **ΔΔ***G*_*int*_ values calculated for the addition of a single-region design to a double-region design are shown as colored arrows in the bottom face.

DeNovoTIMs show a non linear correlation between **Δ***G* at 25 °C and *T*_*m*_, the most commonly used parameters that describe protein stability (inset in Fig. 3C). A similar trend between **Δ***G* at the temperature where it is a maximum (**Δ***G*_Tmax_) and *T*_*m*_ has also been reported for natural and engineered proteins with different sizes and topologies (*34, 45, 46*). Additionally, the global thermodynamic stability can be conveniently described by the area (from 0 °C to *T*_*m*_) under the stability curve (*A*). The advantage of *A* over **Δ***G* at a given temperature is that *A* integrates the conformational stability in a temperature range (*47*). The relative global stability of DeNovoTIMs (*A*/*A*_DeNovoTIM0_) is also correlated with *T*_*m*_ (Fig. 3C). Notably, for DeNovoTIM6, DeNovoTIM11, and DeNovoTIM12, *A*/*A*_DeNovoTIM0_ is nearly ten-fold higher than for DeNovoTIM0 (Fig. 3C and table 1).

The modular strategy used to generate the DeNovoTIMs and the determination of their stabilities allowed us to calculate the contribution of each region to global stability, and also to evaluate the presence of non-additive effects between different regions of the barrel. Non additive effects were evaluated as **ΔΔ***G*_*int*_ through an approach based on thermodynamic double mutant cycles (see Experimental Section). **ΔΔ***G*_*int*_ is also referred to as coupling energy, non-additive effect, interaction energy, and more recently epistatic effect (*48*). Thermodynamic cycles were constructed using the experimental **Δ***G*_25°C_ values obtained from chemical unfolding experiments and linking single-region/double-region designs, and then double-region/triple-region designs as indicated in Fig. S14.

It was found that stabilization is non-additive, consequently, the different barrel regions are coupled, indicating that their contribution to protein stability depends on the structural context. A positive **ΔΔ***G*_*int*_ indicates that the introduction of favorable interactions has a higher stabilizing effect when a nearby region is already mutated. All the **ΔΔ***G*_*int*_ values calculated in Fig. S14 are summarized in the single cube shown in Fig. 3D. **ΔΔ***G*_*int*_ for single and double designs (upper face of the cube) are much smaller than those observed between double- and triple-region designs. The regions that are most energetically coupled in double-region designs are the inner core (DeNovoTIM1) and the upper peripheral core (DeNovoTIM8) (**ΔΔ***G*_*int*_*=* 6.1 kcal mol^-1^, see the upper panel in Fig. S14 and arrows from top to bottom face of the cube in Fig. 3D). Coupling increases considerably when a third region is incorporated on the background of two already mutated regions (**ΔΔ***G*_*int*_*>* 6 kcal mol^-1^, see lower panel in Fig. S14 and arrows on the bottom face of Fig. 3D). The largest **ΔΔ***G*_*int*_ was observed when the DeNovoTIM8 mutations were added to DeNovoTIM11 (**ΔΔ***G*_*int*_*=* 14.2 kcal mol^-1^, see purple arrow in the bottom face of Fig. 3D). Clearly, mutations in one region of the barrel can cause in a non-additive manner the loss or gain of one or more interactions in another distant region of the barrel. The latter indicates that the TIM-barrel fold is suitable for studying modularity and, in general, cooperative effects of proteins. Also, the results presented here suggest that the modular design strategy could be used in the future for the rational stability improvement in other protein topologies.

### Structural analysis of DeNovoTIMs

The structural properties of DeNovoTIMs were examined by X-ray crystallography (table S6). High-resolution data were collected for sTIM11noCys and DeNovoTIM13 (1.88 and 1.64Å, respectively), whereas a low-resolution structure was obtained for DeNovoTIM6 (2.90 Å). All of them showed the designed globular compact TIM-barrel topology (Fig. 4). Structural comparison of the X-ray structures and Rosetta models for sTIM11, sTIM11noCys, DeNovoTIM6, and DeNovoTIM13 showed the lowest RMSD located in the second quarter of the barrel (ranging from 0.27 to 0.68 Å). As previously observed in sTIM11 (*29*), the main structural differences are found in the **α**-helices located at the amino- and carboxyl-terminal ends. In agreement, for all the barrel structures, the RMSD among quarters of the barrel is higher in the first and fourth ones (plot in Fig. 4A). Since the TIM barrel is a closed-repeat protein, contacts between the first and last helices depend on the precise curvature generated by each **α**/**β** unit, therefore geometrical strain may interfere with the proper closure of the barrel.

**Fig. 4.**
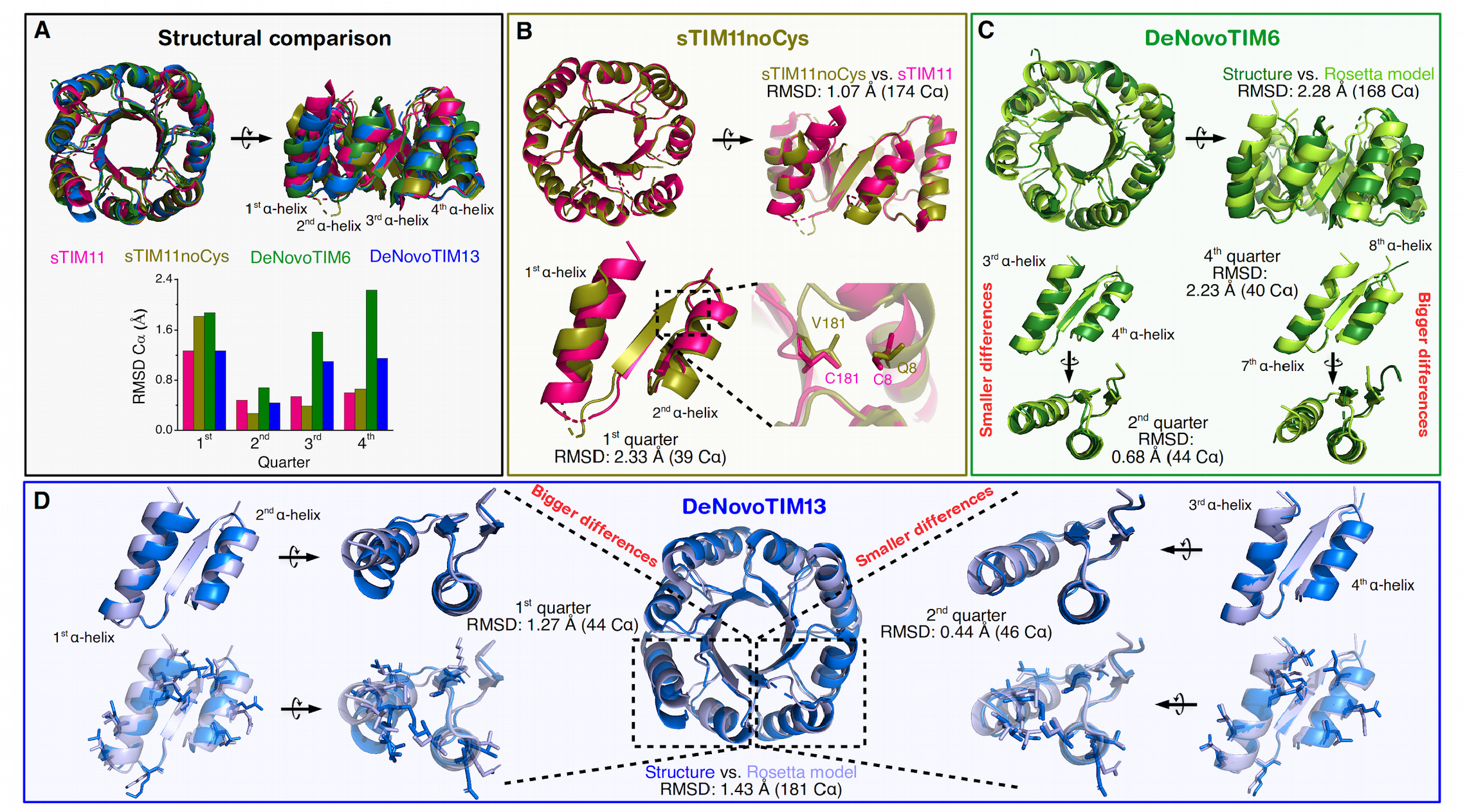
Three-dimensional structures of DeNovoTIMs. **A**) Structural alignment of X-ray structures of sTIM11 (PDB ID: 5BVL), sTIM11noCys (PDB ID: 6YQY), DeNovoTIM6 (PDB ID: 6Z2I), and DeNovoTIM13 (PDB ID: 6YQX). The RMSD C**α** between the structure and the Rosetta model among the quarters in each protein is shown in the lower part of the panel. **B**) Comparison of sTIM11noCys and sTIM11 structures (RMSD: 1.07 Å -174 C**α**-). The mutated residues 8 and 181 in sTIM11noCys are zoomed in the bottom part. **C**) Comparison of the DeNovoTIM6 structure with the Rosetta model (RMSD: 2.28 Å -168 C**α**-). The quarters with the highest and lowest structural similarity are highlighted (bottom left and bottom right, respectively). **D**) Comparison of the DeNovoTIM13 structure with the Rosetta model (RMSD: 1.43 Å -181 C**α**-). The quarters with the highest and lowest structural similarity are highlighted (right and left, respectively). Sidechains of the mutated residues are shown in sticks.

A comparison of the sTIM11noCys and sTIM11 structures showed that removal of the two cysteines causes some structural changes mainly localized in the first and last quarters; the most significant deviations are observed at the amino-terminal region where the first two helices are not well-formed. So even without forming the disulfide bridge, both cysteines in sTIM11 increase the stability and promote a proper closure of the barrel (Fig. 4B and table 1). The other parts of sTIM11noCys adopt almost the same structural arrangement as in sTIM11, except for the **β**_6_/**α**_7_ loop which was not modeled due to an absence of electron density in that region. Thus, although removing the cysteines has effects on stability and structure, sTIM11noCys maintains the general architecture corresponding to the expected TIM barrel.

The thermodynamic properties of DeNovoTIM6 are very similar to those expected for a natural protein (table 1). Unfortunately, due to the low quality of the crystals and therefore the low resolution obtained (2.90 Å), details such as side-chain conformations are not well resolved in the DeNovoTIM6 structure. Nevertheless, it could be verified that the protein is well folded into a compact TIM-barrel (Fig. 4C). As aforementioned for sTIM11 and sTIM11noCys, when the similarity between the structure and the Rosetta model is analyzed, the four quarters in DeNovoTIM6 show different RMSD values (Fig. 4C). The most similar quarter is located in the second region of DeNovoTIM6, whereas the main deviations are located in the first and last quarter of the barrel. Almost all **α**/**β** loops of the barrel are well defined and correspond to the model. However, for some residues within 5 of the 7 **β**/**α** loops no electron density was observed. The main differences observed in the structural analysis between the Rosetta model and the DeNovoTIM6 structure (table S7) are likely due to the low resolution of the data where some residues and side chains are missing in the electron density map. In general, the DeNovoTIM6 structure has high B factors which may reflect higher disorder in the protein crystal or increased flexibility, similar to observations in some regions of sTIM11, namely the amino- and carboxyl-terminal **α**-helices. This could also explain difficulties in obtaining crystals that diffract at higher resolution despite many efforts (see Experimental Section).

As observed in all DeNovoTIMs, the similarities between the DeNovoTIM13 structure and the Rosetta model vary among the four quarters of the barrel (Fig. 4A). The second, third, and fourth quarters display minor differences between the structure and the Rosetta model, with the secondary structure elements and side chains superposing very well. The highest deviations are located at the amino-terminal region that closes the barrel (Fig. 4D). For DeNovoTIM13, the resolution of the crystal structure (1.64 Å) allowed a more in-depth analysis. Most of the hydrogen bonds and salt bridges designed are observed in the DeNovoTIM13 structure. As a consequence of the design strategy, this number is lower than the ones for sTIM11 and sTIM11noCys, stabilizing polar interactions being replaced by an increase in hydrophobic interactions in the DeNovoTIM series. For example, in going from sTIM11 to DeNovoTIM13, a 60 % increase in the total area in hydrophobic clusters was found (3765 vs. 6148 Å^2^); most of this change comes from a three-fold increase in the area of the major hydrophobic cluster (1116 vs. 4351 Å^2^). For DeNovoTIM13, both the area in the major hydrophobic cluster and the total hydrophobic area found in the structure are very similar to those designed (96 and 98 %, respectively; table S7).

One of the main proposed mechanisms for the stabilization of thermophilic proteins is an increase in the number of stabilizing interactions such as salt bridges and hydrogen-bond networks (*5*). In fact, in going from sTIM11 to DeNovoTIM0, a decrease in the number of electrostatic interactions is accompanied by a decrease in stability. In contrast, the structural analysis of DeNovoTIMs showed that these interactions are not clearly related to the observed changes in stability. For example, some of the designs that contained the highest number of polar stabilizing interactions (such as DeNovoTIM1 and DeNovoTIM8) were not the most stable ones, whereas some of the most stable designs (such as DeNovoTIM6 and DeNovoTIMs 12-14) showed a reduction in this type of interaction (table S7). On the contrary, the stability of DeNovoTIMs increases with the number of hydrophobic interactions. The total area, as well as the number of residues and contacts in hydrophobic clusters, are substantially increased in the best first-round designs along with the more stable second- and third-round designs (Fig. S15 and table S7). As discussed in more detail below, this suggests that repacking of the hydrophobic cores is one of the main mechanisms to increase the thermodynamic stability of DeNovoTIMs.

### Epistasis on the stability landscape of *de novo* TIM barrels

To correlate the most common and informative parameters obtained from both temperature and chemical unfolding, *T*_*m*_, **Δ***H*, and **Δ***G*_25°C_ were mapped onto a “stability landscape”, a spatial representation of the observed combinations of these thermodynamic data (Fig. 5 and Experimental Section). Since DeNovoTIMs have different *T*_*m*_ values, experimental **Δ***H* from DSC experiments can not be directly compared. To put the thermodynamic parameters on a similar ground for comparison, **Δ***H* at 85 °C (**Δ***H*_*85°C*_), the average *T*_*m*_ of the DeNovoTIM collection, was calculated using **Δ***H* and **Δ***C*_*P*_ from DSC experiments (table 1).

**Fig. 5.**
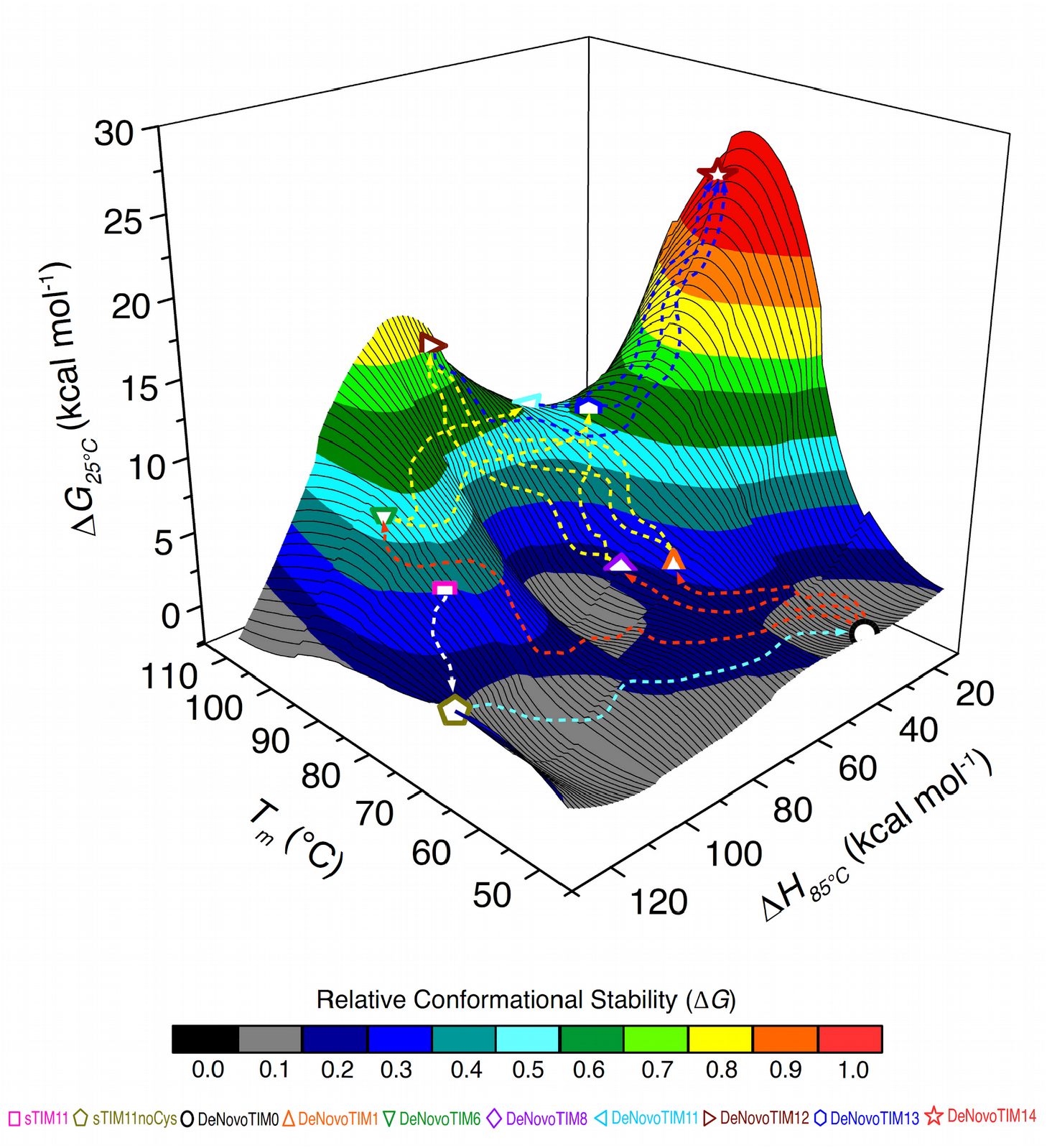
Stability landscape of *de novo* TIM barrels. The stability surface is colored according to normalized **Δ***G*_*25°C*_ values in 0.1 bins. Colored lines represent a possible pathway from one design to another and were drawn only as a guide to the eye: white from sTIM11 to sTIM11noCys, cyan from sTIM11noCys to DeNovoTIM0, orange from DeNovoTIM0 to the first-round DeNovoTIMs, yellow from single-region designs to the double-region ones, and blue from double-region designs to the triple-region design. *T*_*m*_ and **Δ***H* data were obtained from thermal unfolding, whereas **Δ***G* values derive from chemical unfolding. **Δ***H*_*85°C*_ was calculated as indicated in the experimental section, except for DeNovoTIM13 and DeNovoTIM14, where **Δ***H* at *T*_*m*_ was plotted.

The *T*_*m*_ range found in DeNovoTIMs is widely distributed, covering from 47 °C to 109 °C, a range of more than 60 °C in thermostability. Besides, it was possible to design TIM barrels with stabilizing mutations that led to huge differences in stability, even higher than other systems previously reported. The stability landscape of DeNovoTIMs can be compared to that constructed for natural proteins (Fig. S16). The latter is rough, with some regions more populated than others, and explores an ample space due to the diversity in size, topology, oligomeric state, function, and evolutionary history of the variety of natural proteins so far characterized. Interestingly, the comparison shows that several DeNovoTIMs are located in a region of the stability landscape corresponding to low **Δ***H* and high **Δ***G*_25°C_ values, which is not populated, as far as we know, by natural proteins. The modular strategy followed in the DeNovoTIM design rounds can be mapped in this stability landscape. sTIM11, sTIM11noCys, DeNovoTIM0, and most of the first-round designs cover a vast region of the landscape valley, whereas second-round designs are located in a higher stability region. Finally, the third-round design climbs to the highest region of the landscape (Fig. 5).

Assuming additivity, the expected change in stability calculated for DeNovoTIM14 would be the sum of the individual stabilizations provided by all the single-region designs (DeNovoTIM1 + DeNovoTIM6 + DeNovoTIM8) giving a value of 11.2 kcal mol^-1^. However, the stability of DeNovoTIM14 is 23.6 kcal mol^-1^, indicating that more than half of the stabilization comes from positive non-additive effects. The thermodynamic cube presented in Fig. 3D shows that the **ΔΔ***G*_*int*_ mentioned above increases in going from the first-to the second- and third-round designs. Non-additive effects or interaction energies may be referred to as epistasis, a concept traditionally used in genetics to describe the phenotype dependency of a mutation on the genetic state at other sites (*48-50*). Previous studies have explored and analyzed the mechanisms of epistasis within proteins, especially regarding their implications for protein function, evolution, and stability (*51-55*).

Rearrangements in the TIM barrel can influence local changes in other parts of the protein, and these epistatic effects are quantified in the **ΔΔ***G*_*int*_ values whose magnitude for DeNovoTIMs is considerable. The structural analyses suggest that one of the molecular basis of the epistatic effect observed in DeNovoTIMs is likely related to the extension of the hydrophobic cores, particularly to the increase of the major hydrophobic cluster located in the interface between the inner **β**-barrel and the outer **α**-helices (Fig. S15 and table S7). From the first-to the second-round designs, the highest area in hydrophobic clusters was found for DeNovoTIM12, and this corresponds to the highest positive epistatic effect in this round (**ΔΔ***G*_*int*_= 6.1 kcal mol^-1^), whereas the decrease of the hydrophobic cluster area in DeNovoTIM11 (compared to DeNovoTIM1 and DeNovoTIM6) correlates with a negative **ΔΔ***G*_*int*_= -1.8 kcal mol^-1^. From the second-to the third-round designs, the most notable change in hydrophobic area is observed in going from DeNovoTIM11 to DeNovoTIM14, resulting in the highest positive epistatic effect (**ΔΔ***G*_*int*_= 14.2 kcal mol^-1^). The relevance and magnitude of the epistatic or non-additive effects found in DeNovoTIMs, as well as those observed in other reports, suggest that modeling such interactions can improve the success in protein design and engineering.

## Conclusions

Design requires a deep understanding of the relationship between sequence, structure, and stability, and therefore, the combination of thermodynamic and structural data is fundamental to achieve this goal. Here, we designed a family of stable TIM barrels and explored their stability landscape. The TIM-barrel collection reported in this work exhibits a considerable range in thermostability (more than 60 degrees in *T*_*m*_) and conformational stability at 25 °C (more than 22 kcal mol^-1^ in **Δ***G*). These data can now be used to accelerate the development of future custom design protein stability curves which, in turn, will expand the biomedical and biotechnological applications of *de novo* proteins. For example, by fusion to another *de novo* protein, one of the stabilized scaffolds reported here (DeNovoTIM13) has been successfully used to create a reaction chamber on the top of the barrel (56), confirming the convenience of working with robust and stable TIM barrels in the path towards functional *de novo* proteins.

In the same way that one explores the sequence space by studying homologous proteins from different organisms, *de novo* design with a fixed backbone follows a similar strategy generating new sequences within the same topology. It is well known that highly stable proteins can be generated by computational design. However, one of the unexpected findings resulting from the thermodynamic characterization of this family of DeNovoTIMs is that very stable proteins can be obtained in unexplored regions of the stability landscape. The paths followed in the stability landscape of DeNovoTIMs are severely influenced by epistatic effects that appear to arise from an increase in hydrophobic clusters. The design and characterization of stable *de novo* proteins is an essential step on the route to the next generation of new protein functions charting novel sequence space.

## Acknowledgments

We acknowledge financial support and allocation of beamtime by PSI and HZB. We thank the beamline staff at the SLS and at BESSY for assistance, and LANEM-IQ-UNAM for the support in crystal characterization. We thank María Isabel Velázquez López, Laura Iliana Alvarez Añorve, Alma Jessica Díaz Salazar, and Georgina Espinosa Pérez for their competent technical support, Gregor Wiese for generating and crystallizing sTIM11noCys, Noelia Ferruz-Capapey for her help in the structural analyses, as well as Po-Ssu Huang for his comments on the manuscript. We kindly thank all the members of the Fernandez-Velasco, Höcker, and Baker Labs for their constructive suggestions to improve the research.

## Author Contributions

S.R.R., M.C., D.B., B.H., and D.A.F.V. designed the research. D-A.S.M. wrote program code. S.R.R., E.R., C.T., and M.C. collected thermodynamic data. S.R.R., S.K., S.S., Y.G., and A.R.R. solved the crystal structures. S.R.R., M.C., and D.A.F.V. analyzed all the experimental data. S.R.R., B.H., and D.A.F.V. wrote the manuscript. All authors discussed and commented on the manuscript.

## Data and materials availability

All data to support the conclusions of this manuscript are included in the main text and supporting information. Coordinates and structure files have been deposited to the Protein Data Bank (PDB) with accession codes: 6YQY (sTIM11noCys), 6Z2I (DeNovoTIM6), and 6YQX (DeNovoTIM13).

## Supporting Information

Supporting Text, Experimental Section, Supplementary References, Supplementary Figures S1 to S16, and Supplementary Tables S1 to S7 are available in the supporting information of the paper.

