## Supplementary Information for "The stability landscape of *de novo* TIM barrels explored by a modular design approach"

This file includes:

- Supporting Text.
- Experimental Section.
- Supplementary References (1-39).
- Supplementary Figures S1 to S16.
- Supplementary Tables S1 to S7.

### Supporting text

#### Exploratory characterization of first-round designs

The proteins from the first design round (DeNovoTIMs 1-10) as well as DeNovoTIM0, sTIM11, and sTIM11noCys were characterized by circular dichroism (CD) and differential scanning calorimetry (DSC) (Fig. S3). All variants presented far-UV CD spectra with a high content of regular secondary structural elements compatible with α/β proteins. For sTIM11 and sTIM11noCys, the most pronounced minimum of the spectra was observed at 222 nm whereas, for DeNovoTIM0 and DeNovoTIM1-10, it was found at 206 and 208 nm, respectively (Fig. S3 and Fig. S4A). According to the Gibbs Helmholtz equation, high $T_m$ and $\Delta H$ values are reflected in larger areas under the stability curve, therefore, selection of the best designs was based on their thermal unfolding parameters.

Analysis of the internal core designs showed that DeNovoTIM2 and DeNovoTIM3 unfold irreversibly and uncooperatively, whereas DeNovoTIM4 unfolds cooperatively with a $T_m$ value which is 15 degrees higher than DeNovoTIM0 (Fig. S3B). However, DeNovoTIM1 unfolds cooperatively with a further increased $T_m$ that is 24 degrees higher than DeNovoTIM0 (Fig. S3B) and its unfolding $\Delta H$ determined by the DSC endotherm is almost twice than that of DeNovoTIM0 (Fig. S3C). Thus, DeNovoTIM1 was chosen as the best design of this group. Amongst the bottom core designs, DeNovoTIM7 did not overexpress soluble, and no significant efforts were made to solubilize it after unfolding/refolding of inclusion bodies. The thermal unfolding of DeNovoTIM5 is less cooperative and the protein unfolds with a $T_m$ slightly higher than DeNovoTIM0. In contrast, DeNovoTIM6 has a $T_m$ that is 45 degrees higher than DeNovoTIM0 and shows a highly cooperative unfolding transition (Fig. S3E-S3F and table 1). Hence, DeNovoTIM6 is the most successful design within this group. Finally, all top core designs (DeNovoTIM8-10) unfold cooperatively; nevertheless, DeNovoTIM8 displays the highest $T_m$ and $\Delta H$ of this group (Fig. S3H-S3I). Therefore, DeNovoTIM8 is considered the best design among the proteins belonging to this group.

#### Spectroscopic characterization of DeNovoTIMs

sTIM11, sTIM11noCys, and all DeNovoTIM variants presented the characteristic far-UV CD spectra observed for α/β proteins (Fig. 2A). Nevertheless, the secondary structure content deconvoluted from the spectra showed variations in the relative amount of helices and strands, particularly for DeNovoTIM0 and DeNovoTIM1, which have the lowest helical content (table S5). For those DeNovoTIMs for which a three-dimensional structure was obtained (see below), the secondary structure content calculated from the structure and the deconvolution of CD spectra correlates adequately (table S5).

The near-UV CD spectra of DeNovoTIMs (Fig. 2B) showed a peak with fine structure between 290 and 295 nm, characteristic for tryptophan residues. A peak between 275-283 nm was also observed in the designs that contain tyrosine residues (table S3). It is known that the shape and intensities of the near-UV CD spectrum depend not only on the number and identity of aromatic residues but also on their environment and three-dimensional position within the protein core (*1*). The spectra observed in Fig. 2B indicate that aromatic residues in the protein are structured with significant differences in the native environment. The latter is in agreement with the intrinsic fluorescence (IF) spectra, where native $\lambda_{max}$ is in the 329-345 nm range (Fig. 2C and Fig. S6). Likewise, the spectral center of mass (SCM) is between 346 and 357 nm, indicating differences in the environment of aromatic residues among native DeNovoTIMs (table S5). The fluorescence properties of DeNovoTIM0 indicate partial exposure of aromatic residues. All first-round designs show a considerable blue-shift in both $\lambda_{max}$ and SCM; this trend continues in the second- and third-round designs and should be attributed to changes in both the number and the solvent exposure of Trp residues (table S3 and table S5).

#### Residual structure in the thermal unfolding of DeNovoTIMs

All DeNovoTIMs, except DeNovoTIM13 and DeNovoTIM14, showed reversibility i.e. samples were heated up to 125 °C and their endotherms were completely recovered in a second heating scan (Fig. S7). All reversible transitions were well fitted to the two-state model (N⇋U) (Fig. S8) and the calorimetric criterion ($\Delta H_{vH}/\Delta H$) was very close to 1, in agreement with a two-state mechanism (table 1). DSC experiments performed at different protein concentrations (0.25 to 2.5 mg mL$^{-1}$) exhibited the same $T_m$, indicating that thermal unfolding is under thermodynamic control (Fig. S8).

The observed unfolding $\Delta H$ varies greatly, showing values both lower and higher than DeNovoTIM0, sTIM11noCys, and sTIM11 (table 1). When proteins of the same size are compared, the main reasons for finding differences in their unfolding $\Delta H$ are the number of disrupted internal interactions, as well as the hydration of groups exposed upon unfolding (*2*). Extreme scenarios were observed in DeNovoTIMs, e.g. DeNovoTIM6 has a $\Delta H$= 124.9 ± 1.5 kcal mol$^{-1}$, which is 100 kcal mol$^{-1}$ more than that of DeNovoTIM0, and close to the expected value for a protein of 184 residues according to previously reported parametric equations (128.4 ± 3.5 kcal mol$^{-1}$; *3*).

Otherwise, several DeNovoTIMs exhibit small $\Delta H$ values, particularly DeNovoTIM14 ($\Delta H$= 14 kcal mol$^{-1}$; Fig. S10E and table 1). There are at least two main reasons for such a small $\Delta H$, namely either the native state is not fully folded or the unfolded state is not completely unfolded. The spectroscopic, structural, and chemical unfolding properties shown in Fig. 2 indicate that the proteins are well folded in the native state, a conclusion that is supported by the crystal structures (see below). It is known that both $\Delta H$ and $T_m$ decrease in the presence of chemical denaturants (*4*). When DeNovoTIM14 was unfolded by temperature in the presence of urea, the second transition disappeared and the $T_m$ of the observed endotherm remained almost unchanged at ~90 °C (Fig. 2F, Fig. S10B, and Fig. S10D). Unexpectedly, the unfolding enthalpy increased linearly from the value observed without denaturant to a value close to the parametric one at 6.0 M urea (Fig. S10A and Fig. S10E). This atypical behavior in DeNovoTIM14 may be explained by an increased exposure of nonpolar residues upon unfolding in the presence of urea. These results suggest that the reason for the low unfolding enthalpy in some DeNovoTIMs is likely the high content of residual structure in the unfolded state.

Likewise, it has been shown that the residual structure of the unfolded state leads to decreased $\Delta C_P$ values (*5*). For sTIM11, sTIM11noCys, and DeNovoTIM6, the $\Delta C_P$ were close to the estimated value from parametric equations for a protein of that size (2.6 kcal $mol^{-1}$ $K^{-1}$), whereas the other DeNovoTIMs showed a much lower $\Delta C_P$, with DeNovoTIM0 having the lowest value (0.44 kcal $mol^{-1}$ $K^{-1}$, table 1). Accuracy in thermodynamic parameters of unfolding has been discussed over the years and is especially crucial for $\Delta C_P$ due to the baseline shift and its small magnitude. For DeNovoTIMs, the obtention of $\Delta C_P$ was addressed with multiple determinations at varying protein concentration. The standard deviation ranged from 3 to 20 %, very similar to previous estimates for uncertainties in $\Delta C_P$ (*3, 6*). A decrease in unfolding $\Delta C_P$ suggests a non-fully solvated random-coil conformation with residual hydrophobic clusters in the unfolded state. Even though it was not possible to obtain the CD spectra of DeNovoTIMs that display $T_m$ values higher than 90 °C, CD spectra of the unfolded state of the low $T_m$ variants DeNovoTIM0, DeNovoTIM1, and DeNovoTIM8, as well as sTIM11 and sTIM11noCys, clearly showed residual structure (Fig. S4B).

#### Irreversible thermal unfolding of DeNovoTIM13 and DeNovoTIM14

DeNovoTIM13 shows a $T_m$ scan-rate dependent, indicating that its thermal unfolding is under kinetic control (Fig. S9). Because the irreversible thermal unfolding of DeNovoTIM13 and DeNovoTIM14 follows an irreversible two-state mechanism (N→F), well described by a first-order rate constant (*7, 8*), it was possible to determine the activation energy ($E_{act}$) between the native and the transition state. For DeNovoTIM13, the average value obtained from the fitting of each endotherm ($E_{act}$= 118.2 ± 2.7 kcal $mol^{-1}$; Fig. S9A) is within the range reported for natural proteins of similar size. This value agrees with that determined from the Arrhenius plot ($E_{act}$= 118.4 ± 2.4 kcal $mol^{-1}$; Fig. S9B) and with the calculation from the effect of the scan rate on $T_m$ ($E_{act}$= 124.2 ± 1.7 kcal $mol^{-1}$; Fig. S9C).

In contrast, for DeNovoTIM14 a much lower kinetic stability was observed fitting the endotherms to the irreversible two-state model ($E_{act}$= 37.5 ± 0.6 kcal $mol^{-1}$; Fig. S10A), determined from the Arrhenius plot ($E_{act}$= 37.5 ± 0.3 kcal $mol^{-1}$; Fig. S10C), and extrapolating to 0 M urea ($E_{act}$= 37.2 ± 0.8 kcal $mol^{-1}$; Fig. S10F). As a consequence of having very different kinetic stabilities, the estimated half-life of DeNovoTIM13 at 25 °C is 1.8 x $10^{10}$ years and about 196 days for DeNovoTIM14. Clearly, it would be interesting to determine folding/unfolding rates in future kinetic studies of the *de novo* designed TIM barrels.

### Experimental Section

#### Enzymes and biochemicals

All reagents were of analytical grade from Merck KGaA®. Water was distilled and deionized.

#### Design protocol

*De novo* TIM barrels were designed using the Rosetta software suite v.3.2 (*9, 10*; https://www.rosettacommons.org/). All DeNovoTIMs were designed using DeNovoTIM0 as a template. The script used for the DeNovoTIM collection follows and executes the steps indicated by the algorithm as indicated in Fig. S1. In general, the algorithm first selects the symmetry with which it designs the proteins. Once the two-fold or four-fold symmetry was chosen, it selects the number of residues that mutate (depending on whether it is a quarter or half of the protein). Then, an energy minimization step is performed by simulated annealing considering and evaluating the packing and RMSD. Subsequently, it performs a Monte Carlo (MC) fast layer design to improve the packing of the protein's hydrophobic core (in one or several cavities selected according to the regions described in Fig. 1), minimizes the constraints of the main and side chains, and compares each of them with the starting design. For each step, it verifies the RMSD value between both proteins (in the case of DeNovoTIMs, the design was done with a fixed backbone and a cut-off pointed out <0.7 Å). Then, the algorithm filters the results to keep those designs that, with the suggested mutations, were able to increase the packing and preserve the reference topology (ScoreRes: ≤ -1.9, Talaris: ≤ -3.5 BetaNov, Sspred: ≥ 0.85, Packstat: ≥ 0.65). To evaluate if the suggested protein folds as expected, selected designs were computationally validated by a later step of forward folding to predict the *ab initio* three-dimensional structure. The selection was done by an energy score, choosing the designs with the lowest energy value and smallest possible RMSD (located at the bottom left when energy score against RMSD is plotted). In all selected DeNovoTIMs, funnel plots were observed. Finally, the designs were analyzed and the candidates for experimental characterization were selected based on energy criteria, a fewer number of mutations, and physicochemical properties of the suggested mutations.

#### Cloning, overexpression, and protein purification

The nucleotide sequence of all DeNovoTIMs was optimized for expression in *Escherichia coli*. The coding genes were synthesized and cloned into the pET29b(+) vector by GeneScript (New Jersey, USA), except sTIM11noCys, which was cloned into pET21b(+). Proteins were overexpressed in *E. coli* strain BL21(DE3) (Invitrogen®) in 1 L of Terrific Broth (TB) medium supplemented with 30 µg mL$^{-1}$ kanamycin or 100 µg mL$^{-1}$ ampicillin, inoculated with 5 mL preculture and incubated at 37 °C and 200 rpm. After an $OD_{600}$ of 0.6-0.8 was reached, overexpression was induced by adding 1 mM isopropyl-D-1-thiogalactopyranoside (IPTG); growth was continued for 16 hours at 30 °C. After incubation, cells were harvested by centrifugation (Thermo/SLA-3000®, 15 min, 8000 rpm, 4 °C), pellets resuspended in buffer A: 35 mM sodium phosphate, 300 mM NaCl and 35 mM Imidazole pH 8 (supplemented with 0.2 mM of protease inhibitor Phenylmethylsulfonyl fluoride), lysed by sonication (Cole Parmer Ultrasonic Processor®, 10 cycles in 45 s intervals, 30% pulse, 4 °C), and centrifuged again (Sorvall/SS-34®, 40 min, 16000 rpm, 4 °C). In some cases, to increase the efficiency of lysis, the resuspended cells were incubated with lysozyme (250 μg $mL^{-1}$) at 37 °C for 1 hour before sonication. The purification was performed loading the supernatant onto a HisTrap HP column (5 mL; GE Healthcare Life Sciences®) coupled to an ÄKTA system (GE Healthcare Life Sciences®). The unbound fraction was washed out with 20 column volumes (CV) of buffer A. Bound protein was eluted with a linear gradient of 35-500 mM Imidazole using buffer B: 35 mM sodium phosphate, 300 mM NaCl and 500 mM Imidazole pH 8. The pooled fractions were loaded onto a HiLoad 16/600 Superdex 75 preparative grade column (GE Healthcare Life Sciences®). The proteins were purified using isocratic elution with 1.5 CV of buffer C: 150 mM NaCl, 35 mM sodium phosphate pH 8. The fractions corresponding to the monomeric population were pooled and stored at 4 °C for use in subsequent experiments. It should be noted that at the end of protein purification, all designs contain a polyhistidine-tag in the carboxyl-terminal region. For DeNovoTIM11 and DeNovoTIM14, the following purification variables were modified to increase the yield: 0.1 mM of IPTG for induction at $OD_{600}$ of 0.2-0.3, 30 °C and 6 hours for overexpression, buffer A and B containing 1 M NaCl, and buffer C with 300 mM NaCl. At each step of the purification process, aliquots were taken to quantify the amount of protein and to calculate the corresponding purification tables. The final yields are indicated in table S5.

#### Far- and Near-UV Circular Dichroism

Circular Dichroism (CD) spectra were collected in buffer D: 10 mM sodium phosphate pH 8 in a Chirascan Spectropolarimeter using a Peltier device to control the temperature (Applied Photophysics®). For Far-UV spectra, 0.4 mg $mL^{-1}$ of DeNovoTIM was used for all measurements (1 nm bandwidth, 185-260 nm wavelength range, 1 mm cuvette). For Near-UV spectra, 1 mg $mL^{-1}$ of DeNovoTIM was used for all measurements (1 nm bandwidth, 250-350 nm wavelength range, 10 mm cuvette). The spectra for thermally-unfolded states were collected at 90 °C. Spectra for chemically-unfolded states were collected at 9 M urea for all DeNovoTIMs, except for DeNovoTIM14, which was collected at 7 M GdnHCl. Raw data were converted to mean residue molar ellipticity ($[\theta]$) using: $[\theta] = \theta/(l\ C\ N_r)$, where $\theta$ is ellipticity collected in millidegrees, l is the cell path length in mm, C is the DeNovoTIM molar concentration, and $N_r$ the number of residues per protein. Far-UV spectra were deconvoluted with CDNN (*11*).

#### Intrinsic Fluorescence

Intrinsic Fluorescence (IF) spectra were collected on a PC1 ISS Spectrofluorometer (Champaign IL-USA®) equipped with a Peltier device controlling the temperature. In all measurements, protein concentration was 0.4 mg $mL^{-1}$ in buffer D: 10 mM sodium phosphate pH 8 (1 nm bandwidth slits, 295 nm excitation wavelength, 310–450 nm emission wavelength range). Spectra for chemically-unfolded states were collected at 9 M urea for all DeNovoTIMs, except for DeNovoTIM14, which was collected at 7 M GdnHCl. Fluorescence spectral center of mass (SCM) was calculated from intensity data ($I_\lambda$) obtained at different wavelengths ($\lambda$): SCM= $\sum \lambda I_\lambda / \sum I_\lambda$.

#### Three-dimensional structure determination

DeNovoTIMs were concentrated with Amicon Ultra centrifugal filter units (Millipore®) and dialyzed in buffer C: 10 mM sodium phosphate pH 8, 150 mM NaCl. Sitting-drop vapor-diffusion method, and JCSG Core I-IV, JCSG +, Classics I-II, PACT, PEGs I-II, and $AmSO_4$ screening suites (Qiagen®) were used to screen crystallization conditions in 96 well Intelli plates (Art Robbins Instruments®) stored at 20 °C in the hotel-based Rock Imager RI 182 (Formulatrix®). 0.8 μL drops were prepared in a 1:1 ratio with mother liquid using a nanodispensing crystallization robot Phoenix (Art Robbins Instruments®) and then optimized by multiple crystallization rounds using a sitting-drop vapor-diffusion method. To improve the diffraction quality of DeNovoTIM crystals, different pre- and post-crystallization methods were used: reductive methylation (JBS Methylation Kit, Jena Biosciences®), seeding, additive screening, controlled dehydration, cryoprotection screening, crystal annealing, and room-temperature diffraction. In total, more than 300 different crystals in various conditions were tested.

Suitable crystals for X-ray diffraction were found in the following conditions: sTIM11noCys: 0.2 M Ammonium Sulfate, 0.1 M Trisodium Citrate pH 5.6, 25% w/v Polyethylene glycol (PEG) 4000, with a protein concentration of 15 mg $mL^{-1}$; DeNovoTIM6: 0.095 M Sodium Citrate pH 5.0, 19% v/v Isopropanol, 25% w/v PEG 4000, 5% v/v Glycerol, with a protein concentration of 8.6 mg $mL^{-1}$; DeNovoTIM13: 0.17 M Sodium Acetate, 0.085 M Tris pH 8.5, 25.5% w/v PEG 4000, 15% v/v Glycerol, with a protein concentration of 10 mg $mL^{-1}$.

For sTIM11noCys and DeNovoTIM13, diffraction data were collected at 100 K at the Swiss Light Source at the Paul Scherrer Institute in Villigen (Switzerland) (X10SA-PXII beamline for sTIM11noCys and X06DA-PXIII beamline for DeNovoTIM13) using a wavelength of 1 Å and a PILATUS 6M detector for sTIM11noCys and a PILATUS 2M-F detector for DeNovoTIM13 (*12, 13*). For DeNovoTIM6, diffraction data were collected at 100 K at the Berlin Electron Storage Ring Society for Synchrotron Radiation beamline 14.2 (BESSY BL14.2) operated by Helmholtz-Zentrum Berlin using a wavelength of 0.91 Å and a PILATUS3S 2M detector (*14*).

Diffraction data were processed with the X-ray Detector Software (XDS) using XDSAPP v.2.0 (*15, 16*) for sTIM11noCys and DeNovoTIM13, and DIALS (*17*) for DeNovoTIM6. For data reduction, criteria used to cut off the data were the resolution shell with a meanI/sigma(I) between 1-2 and the best CC1/2 according to redundancy and completeness. The structures were solved by molecular replacement with PHASER in the PHENIX software suite v.1.17 (*18*) using sTIM11 (PDB ID: 5BVL) as a starting model for sTIM11noCys and the own Rosetta model for DeNovoTIM6, DeNovoTIM13, and DeNovoTIM14. Refinement was done with phenix.refine (*18*). The model was improved by map inspection and iteratively manual rebuilding performed in COOT v.0.9 (*19*). The final coordinates were validated with PDB_REDO (*20*), MolProbity v.4.2 (*21*), and the Protein Data Bank validation service (*22*); in all servers, the 3D-structure satisfied all quality criteria. The coordinates and structure factors were deposited in the PDB with accession codes: 6YQY (sTIM11noCys), 6Z2I (DeNovoTIM6), and 6YQX (DeNovoTIM13). The figures were created using PyMOL Molecular Graphics System v.4.5.0 (Schrodinger, LLC).

#### Analytical Size Exclusion Chromatography

Hydrodynamic measurements were performed on a Superdex 75 10/300 GL analytical column coupled to an ÄKTA System (GE Healthcare Life Sciences®). All experiments were performed in buffer C: 10 mM sodium phosphate pH 8, 150 mM NaCl at 25 °C and a protein concentration range from 0.01 to 2.0 mg mL$^{-1}$. Experimental molecular weight, Stokes-radii, and oligomeric state were calculated from elution volumes and a calibration curve derived from 7 different known proteins.

#### Thermal unfolding followed by Circular Dichroism

Temperature-induced unfolding was monitored by CD at 222 nm as a function of temperature using 0.4 mg mL$^{-1}$ in buffer D: 10 mM sodium phosphate pH 8, a heating rate of 1.0 and 1.5 K min$^{-1}$, and a 1 mm path-length cell. The changes in the CD signal were normalized to the fraction of unfolded molecules ($f_U$) by:

$$f_U = \frac{y_{obs} - (y_N + m_N T)}{(y_U + m_U T) - (y_N + m_N T)} \quad \text{(Eqn. 1)}$$

where $y_{obs}$ is the experimentally observed CD signal at a given temperature, and ($y_N$+$m_N$T) and ($y_U$+$m_U$T) are the linear fitting equations corresponding to the native and unfolded regions, respectively. $T_m$ values were estimated from normalized data fitted with a Boltzmann-type function:

$$f_U = \frac{-1}{\left(1 + e^{\frac{T - T_m}{a}}\right)} + 1 \quad \text{(Eqn. 2)}$$

where $a$ is related to the slope of the transition.

#### Thermal unfolding followed by Differential Scanning Calorimetry

Differential Scanning Calorimetry (DSC) scans were carried out in a VP-Capillary DSC system (MicroCal®, Malvern Panalytical). Samples were prepared by exhaustive dialysis in buffer D: 10 mM sodium phosphate pH 8 and then degassed at room temperature. To ascertain proper instrument equilibration, two buffer-buffer scans were performed before each protein-buffer scan (Fig. S7). Corresponding buffer-buffer traces were subtracted from each endotherm. For all proteins a reheating scan was performed to determine the reversibility or irreversibility of the process (Fig. S7). Reversibility percentage was calculated by comparing the calorimetric Δ*H* (area under the curve) recovered in the second scan and that obtained in the first one ($\Delta H_{secondscan}/\Delta H_{firstscan}$) *100. To verify that irreversibility was not the result of a too high final scanning temperature, the first scans were also performed heating near the $T_m$. For DeNovoTIMs with a reversible thermal unfolding, protein concentration varied from 0.25 to 2.5 mg $mL^{-1}$ and scan rates from 0.5 to 3.0 K $min^{-1}$, except for DeNovoTIM0 where protein concentration varied from 1 to 5 mg $mL^{-1}$. For DeNovoTIMs with an irreversible thermal unfolding, protein concentration was 1 mg $mL^{-1}$ and scan rates from 0.5 to 3.0 K $min^{-1}$. For DeNovoTIM14 in native conditions, protein concentration was increased to 2.5 and 4.5 mg $mL^{-1}$ to determine accurately the transition. For DeNovoTIM14 in the presence of urea, all the scans were done at 1 mg $mL^{-1}$ from 2.0 to 6.0 M urea with samples incubated for 6 hours at 10 °C. Origin v.9.0 (OriginLab Corporation, Northampton, MA, USA.) with MicroCal software was used for data analysis.

*Thermodynamic parameters from reversible DSC transitions*

DSC endotherms were fitted to equilibrium two-state model (N⇋U):

$$C_P(T) = B_0 + B_1 T + f(T)\Delta C_P + \frac{\Delta H(T)}{R T_m^2}\left[\frac{1-f(T)}{1-n+\frac{n}{f(T)}}\right] \quad \text{(Eqn. 3)}$$

where $B_0$ and $B_1$ are pre- and post-transition constants, n is the number of subunits in the native protein (1 for all DeNovoTIMs) and *f(T)* is the protein fraction in the folded monomeric state, producing Δ*H* (at $T_m$), $\Delta C_P$, and $T_m$. The thermodynamic parameters reported are the average of ten experiments carried out in the 0.25 to 2.5 mg $mL^{-1}$ range. The van't Hoff enthalpy ($\Delta H_{vH}$) was evaluated by (*23*):

$$\Delta H_{vH} = \frac{4 R T_m^2 C_{P,T_m}}{\Delta H} \quad \text{(Eqn. 4)}$$

where *R* is the universal gas constant, $T_m$ is the temperature at which $C_P$ is maximal, $C_{P,Tm}$ is the heat capacity value at $T_m$, and Δ*H* is the total calorimetric enthalpy of the endotherm.

*Thermodynamic parameters from irreversible DSC transitions*

Calorimetric transitions were adequately described by the two-state irreversible model (N→F) where N is the native protein and F is the final state (*7, 8*). The kinetic conversion from N to F is described by a first-order rate constant (*k*) changing with temperature according to the Arrhenius equation:

$$k = \exp\left[\frac{-E_{act}}{R}\left(\frac{1}{T} - \frac{1}{T^{'}}\right)\right] \quad \text{(Eqn. 5)}$$

where $T^*$ is the temperature at which the $k$= 1 min$^{-1}$ and $E_{act}$ is the activation energy between the native and the transition states that describes the unfolding process. The apparent heat capacity is given by:

$$C_P^{APP} = \frac{\Delta H\, E_{act}}{R T_m^2} \exp(x) \exp[-\exp(x)]; x = \frac{E_{act}}{R T_m^2}(T - T_m) \quad \text{(Eqn. 6)}$$

where $T$ is the temperature and $\Delta H$ is the unfolding enthalpy. The $E_{act}$ was also obtained following these two procedures: from the slope of Arrhenius plots, i.e. ln $k$ vs. 1/T; and derived from a data consistency test, evaluating the effect of scanning rate ($\nu$) on $T_m$ (*24*).

#### Chemical-induced unfolding

All experiments were carried out at a protein concentration of 0.1 mg mL$^{-1}$ in buffer D: 10 mM sodium phosphate pH 8 at 25 °C. To determine whether urea induced unfolding was reversible, unfolding and refolding experiments were assayed. For unfolding experiments, native DeNovoTIM was the initial state, whereas for refolding, the starting state was the unfolded DeNovoTIM incubated overnight in 9.0 M urea. Thereafter samples were incubated at different concentrations of urea (0-9.0 M), either increasing or decreasing the initial concentration (for unfolding and refolding experiments, respectively). Intrinsic fluorescence of both, unfolding and refolding samples, was measured at different times to determine the equilibrium time. Once the equilibrium time was found, unfolding experiments with samples incubated for 12 hours and followed by CD and IF were performed as aforementioned. IF data at fixed emission wavelength and CD data at 222 nm were both collected over 2 minutes at each urea concentration. The changes in IF and CD were normalized to the fraction of unfolded molecules ($f_U$) by:

$$f_U = \frac{y_{obs} - (y_N + m_N[urea])}{(y_U + m_U[urea]) - (y_N + m_N[urea])} \quad \text{(Eqn. 7)}$$

where $y_{obs}$ is the experimentally observed IF and CD signal at a given temperature, and ($y_N$+$m_N$[urea]) and ($y_U$+$m_U$[urea]) are the linear fitting equations corresponding to the native and unfolded regions, respectively. All two-state transitions were fitted to Santoro and Bolen equation (*25*) which assumes a two-state model (N⇌D):

$$f_U = \frac{(y_N + m_N[urea]) + (y_U + m_U[urea])\, e^{\frac{-\Delta G^{H2O} - m[urea]}{RT}}}{1 + e^{\frac{-\Delta G^{H2O} - m[urea]}{RT}}} \quad \text{(Eqn. 8)}$$

where $\Delta G^{H2O}$ is the unfolding free energy in absence of denaturant, $m$ is $\Delta G$/[urea], $T$ is the temperature of the experiment (25 °C), and ($y_N$+$m_N$[urea]) and ($y_U$+$m_U$ [urea]) are the linear fitting equations for the pre- and post-transition states. The chemical unfolding transitions for DeNovoTIM14 in GdnHCl were fitted to a three-state model with an intermediate:

$$f_U = \frac{\left(\left(\left(y_U + m_U[GdnHCl]\right)\right) K_1 K_2 + \left(y_N + m_N[GdnHCl]\right) + \left(\left(y_I + m_I[GdnHCl]\right)\right) K_1\right)}{1 + K_1 + K_1 K_2} \quad \text{(Eqn. 9)}$$

where $K_1 = e^{\frac{-\Delta G_{NtoI} - m_{NtoI}[GdnHCl]}{RT}}$, $K_2 = e^{\frac{-\Delta G_{ItoU} - m_{ItoU}[GdnHCl]}{RT}}$, $\Delta G_{NtoI}$ and $\Delta G_{ItoU}$ is the unfolding free energy from native state to intermediate and from intermediate to unfolded state, $m_{NtoI}$ and $m_{ItoU}$ is $\Delta G$/[GdnHCl] of each step, $T$ is the temperature of the experiment (25 °C), and ($y_N$+$m_N$[GdnHCl]), ($y_I$+$m_I$[GdnHCl]), and ($y_U$+$m_U$[GdnHCl]) are the linear fitting equations for native, intermediate, and unfolded states, respectively. Similar $\Delta G$ values were obtained when experimental protein concentration was increased five fold, ruling out the possibility of a bimolecular association/folding step.

#### Stability curve and global thermodynamic stability

Global stability curves, $\Delta G$(T), were calculated using the thermodynamic parameters obtained from DSC experiments and the Gibbs-Helmholtz equation (*26*):

$$\Delta G(T) = \Delta H\left(1 - \frac{T}{T_m}\right) - \Delta C_P\left(T_m - T + T ln\left(\frac{T}{T_m}\right)\right) \quad \text{(Eqn. 10)}$$

The area under the stability curve is a measure of the global stability of the protein (*27*). It was calculated integrating equation 10 from the lowest temperature at which the protein is in the liquid state i.e. 0 °C (273.15 K) to $T_m$:

$$Area = \left(\left(\Delta H - T_m \Delta C_P\right)\left(T_m - T\right)\right) - \left(\frac{\Delta H}{2T_m} - \frac{\Delta C_P}{2}\right)\left(T_m^2 - T^2\right) + \left(\frac{\Delta C_P}{4} T_m^2\right) + \frac{\Delta C_P}{2}\left(T^2 \ln\frac{T}{T_m} - \frac{T^2}{2}\right) \quad \text{(Eqn. 11)}$$

#### Stability landscape

The stability landscape was constructed by plotting $T_m$ and $\Delta H_{85°C}$ obtained from thermal unfolding experiments, and $\Delta G_{25°C}$ obtained from chemical unfolding data. $\Delta H$ at 85 °C ($\Delta H_{85°C}$), the average $T_m$ of the DeNovoTIM collection, was calculated as follows (*28*):

$$\Delta H_{85°C} = \Delta H + \Delta C_P\left(85°C - T_m\right) \quad \text{(Eqn. 12)}$$

where $T_m$, $\Delta H$ and $\Delta C_P$ are the experimental values obtained from DSC experiments for each protein, and 85 °C is the reference temperature. $\Delta H_{85°C}$ was not calculated for DeNovoTIM13 and DeNovoTIM14, because their irreversible thermal unfolding hampered the determination of $\Delta C_P$.

The 3D surface map was calculated using an XYZ gridding approach for randomly spaced data based on the modified Shepard's method. The expanded matrix was a rectangular array with $\Delta G_{25°C}$ as Z values whose columns were mapped to $T_m$ as X values and rows to $\Delta H$ as Y values. The method constructs a function $F(x,y)$ go through the experimental data ($T_m$, $\Delta H$, and $\Delta G_{25°C}$) and interpolates ($F(x_i, y_i) = z_i$) for all irregular distributed points ($x_i$, $y_i$, $z_i$). The stability surface was constructed with the software Origin v.9.0 (OriginLab Corporation, Northampton, MA, USA.) and colored according to normalized $\Delta G_{25°C}$ values in 0.1 bins. It should be noted that although $T_m$, $\Delta H$, and $\Delta G$ are related by equation 10, their surface representation in 3D requires a common $\Delta C_P$. Therefore, the stability surfaces shown in Fig. 5 and Fig. S16 are not a 3D fitting to the Gibbs-Helmholtz equation.

#### Thermodynamic double-mutant cycles

To calculate non-additive effects between different DeNovoTIM barrel regions, an approximation based on double mutant cycles was used (*29-31*). The thermodynamic cycles were constructed using the experimental $\Delta G_{25°C}$ values obtained from chemical unfolding experiments and linking single-region/double-region designs and double-region/triple-region designs as indicated in Fig. S14.

Each corner of the square represents a different DeNovoTIM where the mutations are located in a specific region of the barrel or in a combination of them. For double-region cycles (upper panel), from the first to the second design round, $\Delta G_1$ and $\Delta G_2$ are the changes in stability produced when a single region of the barrel was mutated, $\Delta G_3$ and $\Delta G_4$ are the changes in stability generated when the same mutations are evaluated in the background of another first-round design. In the triple-region cycles (lower panel), from the second to the third design round, $\Delta G_1$ and $\Delta G_3$ are the changes in stability produced when the mutations of a single region are introduced in the background of DeNovoTIM0 or in a double-region design, whereas $\Delta G_2$ and $\Delta G_4$ are the changes in stability generated when a double-region design was incorporated in the background of DeNovoTIM0 or in a single region design, respectively.

Considering that $\Delta G$ is a state property, if two regions of the barrel are energetically-independent, their effects will be additive and not coupled. Therefore, stability changes linked to a particular region will result in the same values on parallel sides of the square, i.e., $\Delta G_1=\Delta G_3$ and $\Delta G_2=\Delta G_4$. Any difference the values on the parallel sides of the squares indicates a deviation from additivity and measures the coupling energy between different regions of the barrel, given by $\Delta\Delta G_{int}= \Delta G_4-\Delta G_2= \Delta G_3-\Delta G_1$, where $\Delta\Delta G_{int}$ values have been referred as coupling energy, non-additive effects, interaction energies, and more recently epistatic effects (*29*).

#### Sequence and structural analysis

Sequence alignment was performed with MAFFT v.7.450 (*32*) using the secondary structure information from the sTIM11 structure (PDB ID: 5BVL). Sequence identity was calculated with the SIAS server (Universidad Complutense de Madrid, 2013). Structural alignments and RMSD calculations were performed using PyMOL Molecular Graphics System v.4.5.0 (Schrodinger, LLC). Cavity volumes were calculated with MOLE v.2.5 (*33*) using a standard probe radius of 5 Å and an interior threshold of 1.1 Å with a non-directed exploration path. The accessible surface area (ASA) was calculated with VADAR v.1.8 (*34*). In these analyses, changes in ASA for the unfolded state were calculated with an extended Gly-X-Gly peptide. Hydrogen bonds, as well as salt bridges, were calculated using HBPLUS v.3.06 (*35*) and ESBRI (*36*) with default parameters for distances and angles. A salt bridge was assigned when two atoms of opposite charge were observed within 4 Å. Hydrophobic clusters (formed by ILV residues) were calculated following an algorithm previously reported by Sobolev (*37*) and available in the *ProteinTools* server developed by Dr. Noelia Ferruz-Capapey from the Höcker Lab (https://proteintools.uni-bayreuth.de/).

#### Supplementary figures

##### List of supplementary figures


- **Fig. S1**: Flowchart of the Rosetta design protocol used to generate the DeNovoTIM collection.
- **Fig. S2**: Sequence alignment of DeNovoTIM barrels.
- **Fig. S3**: Exploratory characterization of first-round designs.
- **Fig. S4**: CD spectra in the peptidic region for DeNovoTIMs.
- **Fig. S5**: CD spectra in the aromatic region for DeNovoTIMs.
- **Fig. S6**: Intrinsic fluorescence spectra of DeNovoTIMs.
- **Fig. S7**: DSC instrument equilibration and thermal unfolding reversibility assessment of DeNovoTIMs.
- **Fig. S8**: DSC endotherms of DeNovoTIMs.
- **Fig. S9**: Irreversible thermal unfolding of DeNovoTIM13.
- **Fig. S10**: Irreversible thermal unfolding of DeNovoTIM14 in the presence of urea.
- **Fig. S11**: $\Delta C_P$ determination of DeNovoTIMs at different protein concentrations.
- **Fig. S12**: Chemical unfolding of DeNovoTIMs followed by CD and IF (raw data).
- **Fig. S13**: Chemical unfolding of DeNovoTIMs followed by CD and IF (normalized data).
- **Fig. S14**: Thermodynamic cycles for DeNovoTIMs.
- **Fig. S15**: Hydrophobic clusters of DeNovoTIMs.
- **Fig. S16**: Stability landscape of natural proteins.

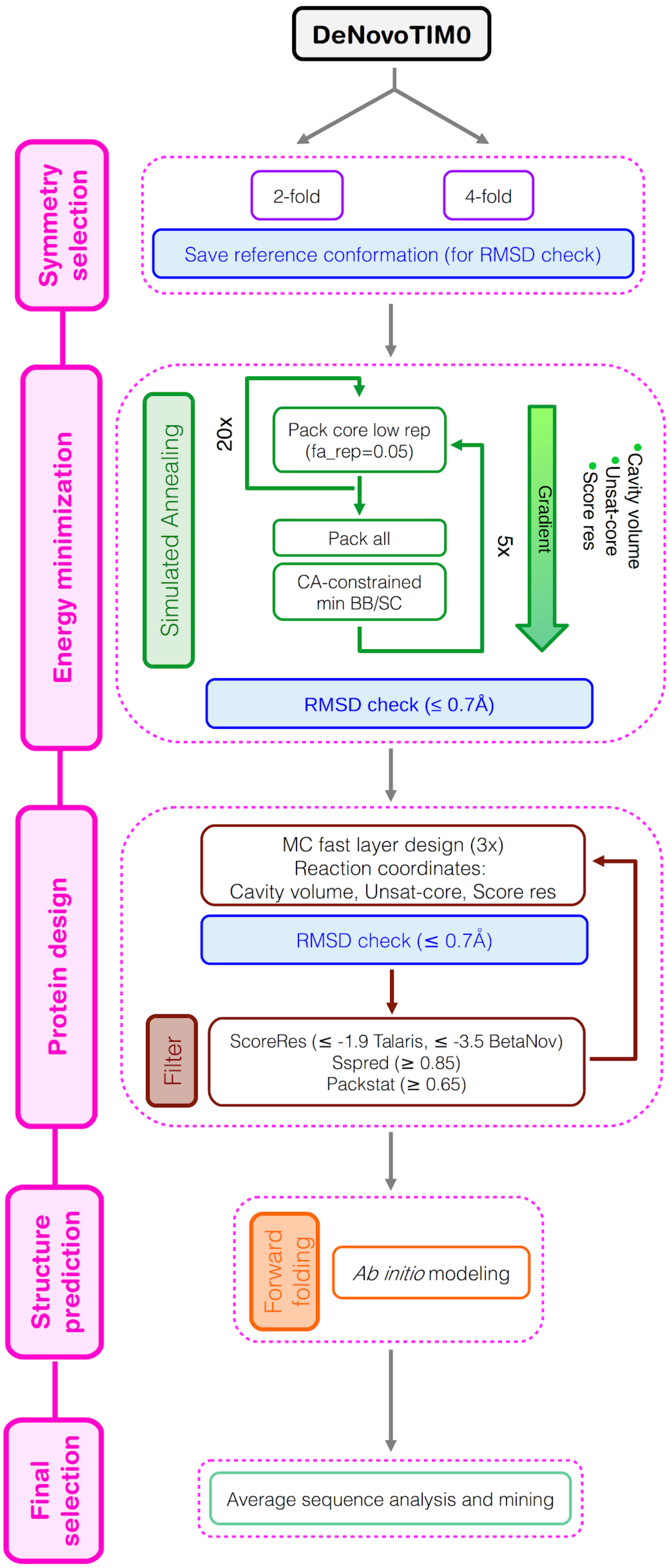


Fig. S1. Flowchart of the Rosetta design protocol used to generate the DeNovoTIM collection.

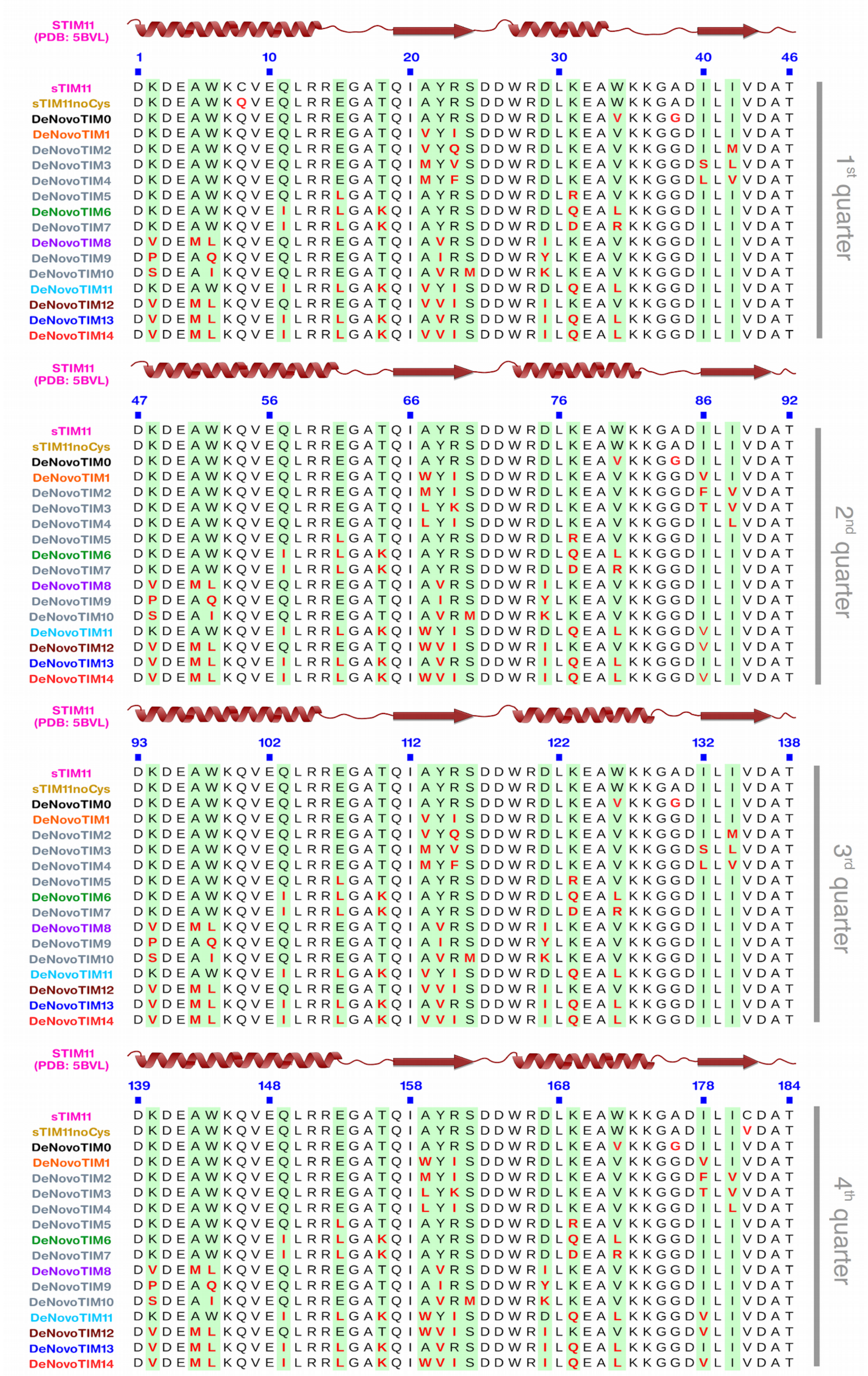


**Fig. S2. Sequence alignment of DeNovoTIM barrels**. On top of each TIM barrel quarter, the secondary structure of sTIM11 is shown (PDB ID: 5BVL). Mutations incorporated in each design are highlighted in red. sTIM11noCys is the symmetric version of sTIM11 removing the cysteine residues (C8Q/C181V). DeNovoTIM0 contains the additional mutations W34V and A38G, as well as their symmetry-related positions. See table S2 for a complete list of the mutations.

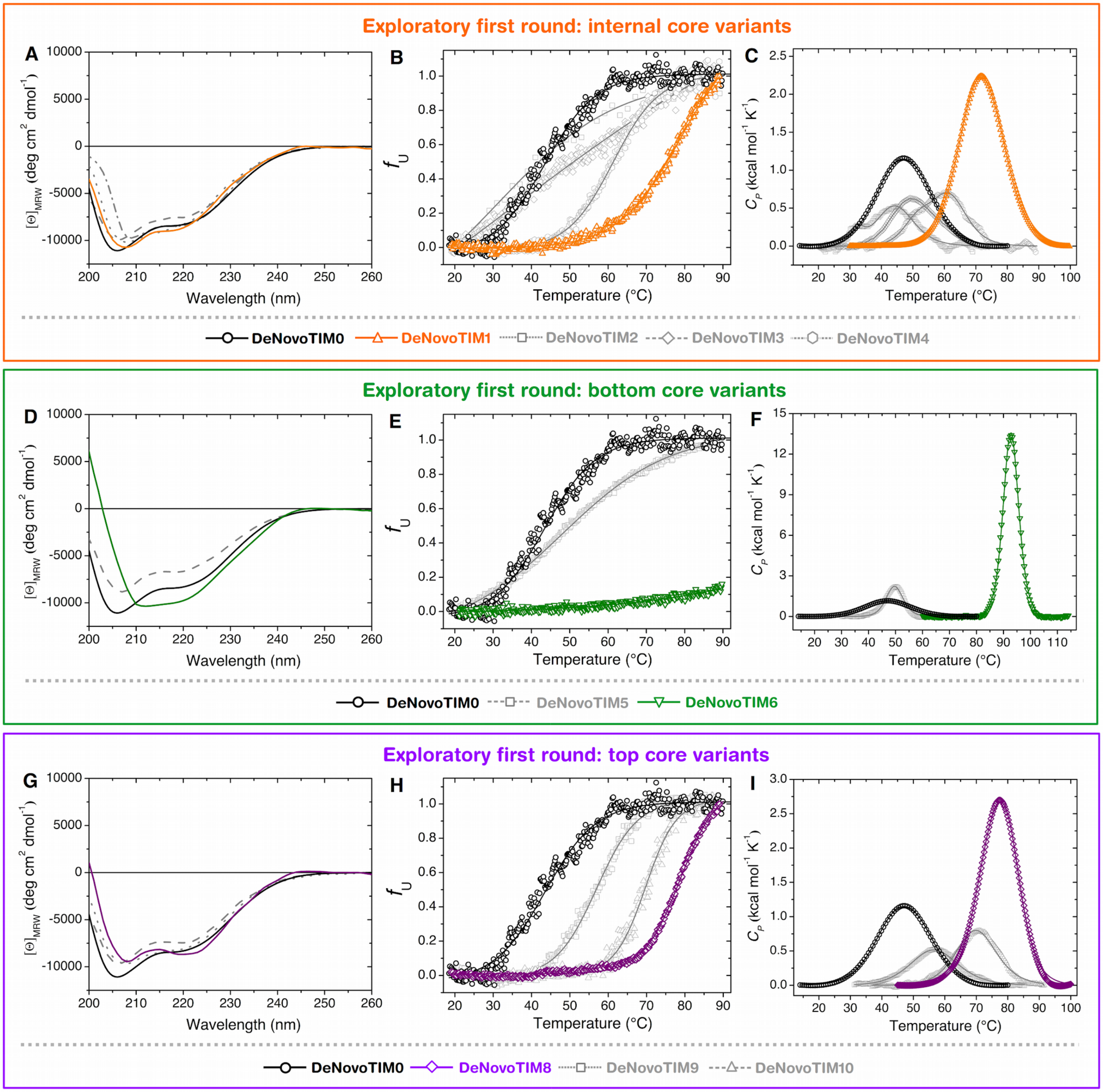


**Fig. S3. Exploratory characterization of first-round designs**. Far-UV CD spectra are shown in panels **A**, **D**, and **G**. Normalized thermal unfolding data followed by $CD_{222\ nm}$ are presented in panels **B**, **E**, and **H**. Thermal unfolding experiments followed by DSC are shown in panels **C**, **F**, and **I**. Selected variants of each design group are highlighted in orange (DeNovoTIM1), green (DeNovoTIM6), or purple (DeNovoTIM8). In all experiments protein concentration was 0.4 mg $mL^{-1}$ in 10 mM sodium phosphate pH 8.

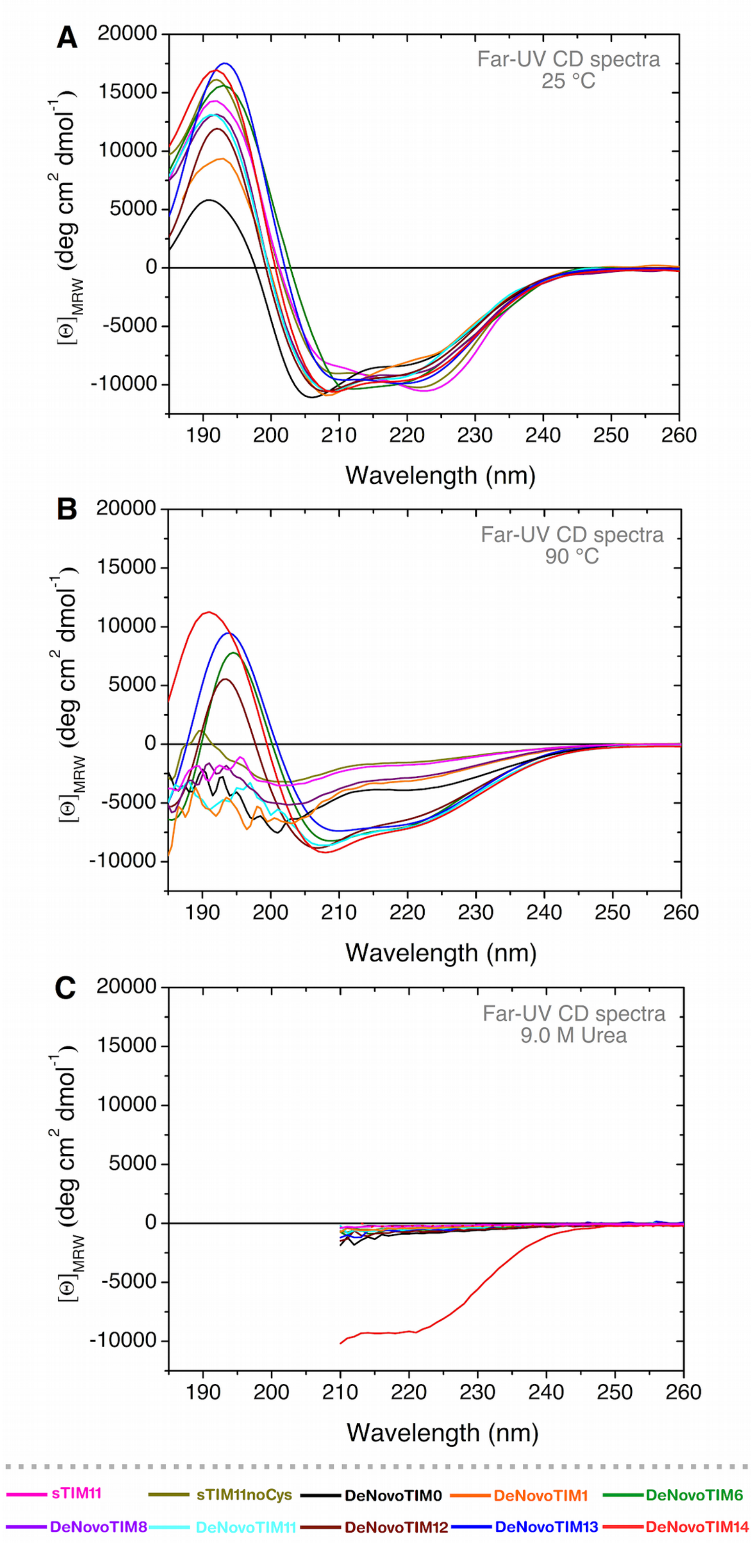


**Fig. S4. CD spectra in the peptidic region for DeNovoTIMs.** Far-UV CD spectra at 25 °C (**A**), 90 °C (**B**), and 9.0 M urea (**C**). Note that for DeNovoTIM14 at 9.0 M urea (solid red line) the spectrum is very similar to the native one (panel **A**). In 7.0 M GdnHCl the protein is completely unfolded and its spectrum is identical to all other DeNovoTIMs unfolded in 9.0 M urea (dotted red line). In all experiments protein concentration was 0.4 mg mL$^{-1}$ in 10 mM sodium phosphate pH 8.

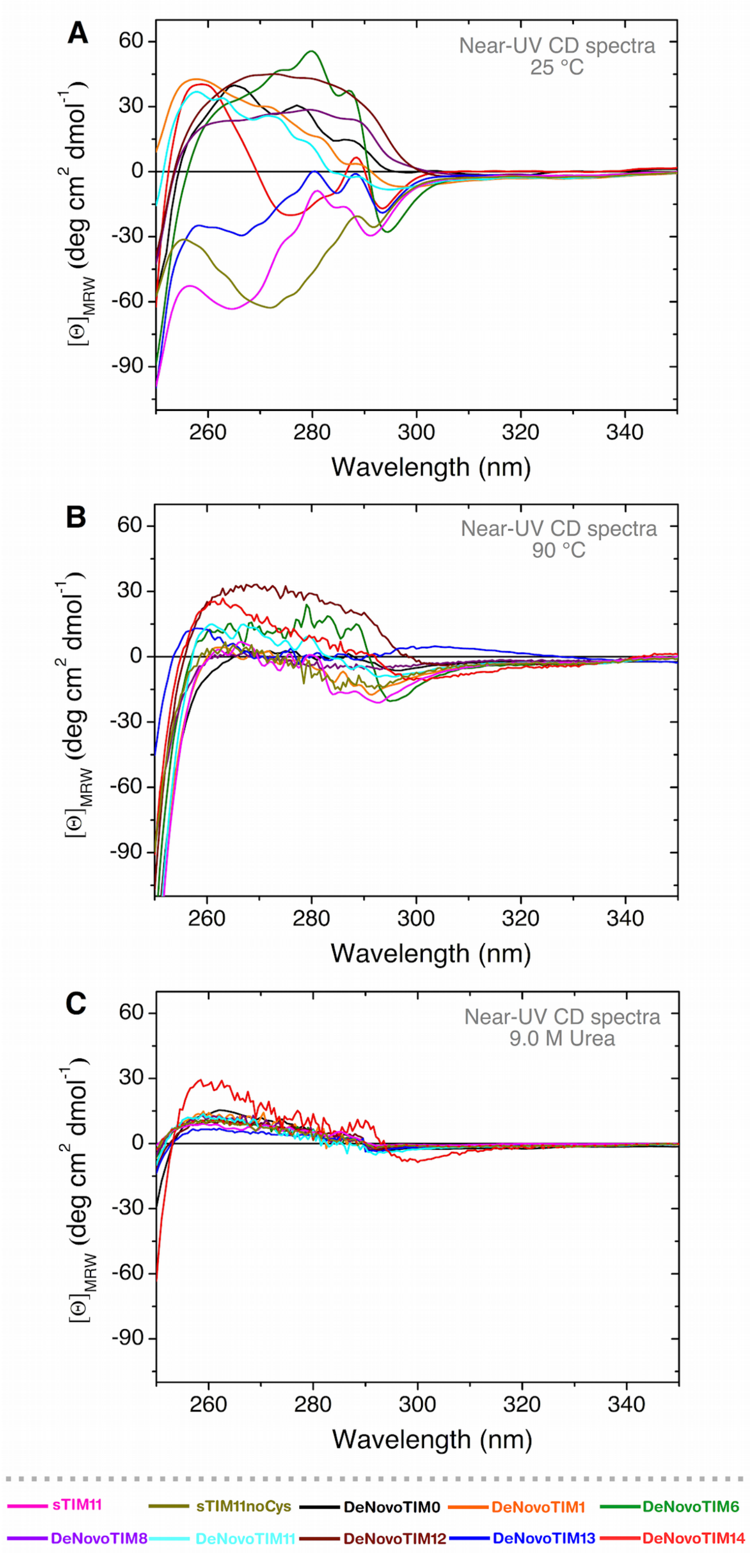


**Fig. S5. CD spectra in the aromatic region for DeNovoTIMs.** Near-UV CD spectra at 25 °C (**A**), 90 °C (**B**), and 9.0 M urea (**C**). Note that for DeNovoTIM14 at 9.0 M urea (solid red line) the spectrum is very similar to the native one (panel **A**). In 7.0 M GdnHCl the protein is completely unfolded and its spectrum is identical to all other DeNovoTIMs unfolded in 9.0 M urea (dotted red line). In all experiments protein concentration was 0.4 mg mL$^{-1}$ in 10 mM sodium phosphate pH 8.

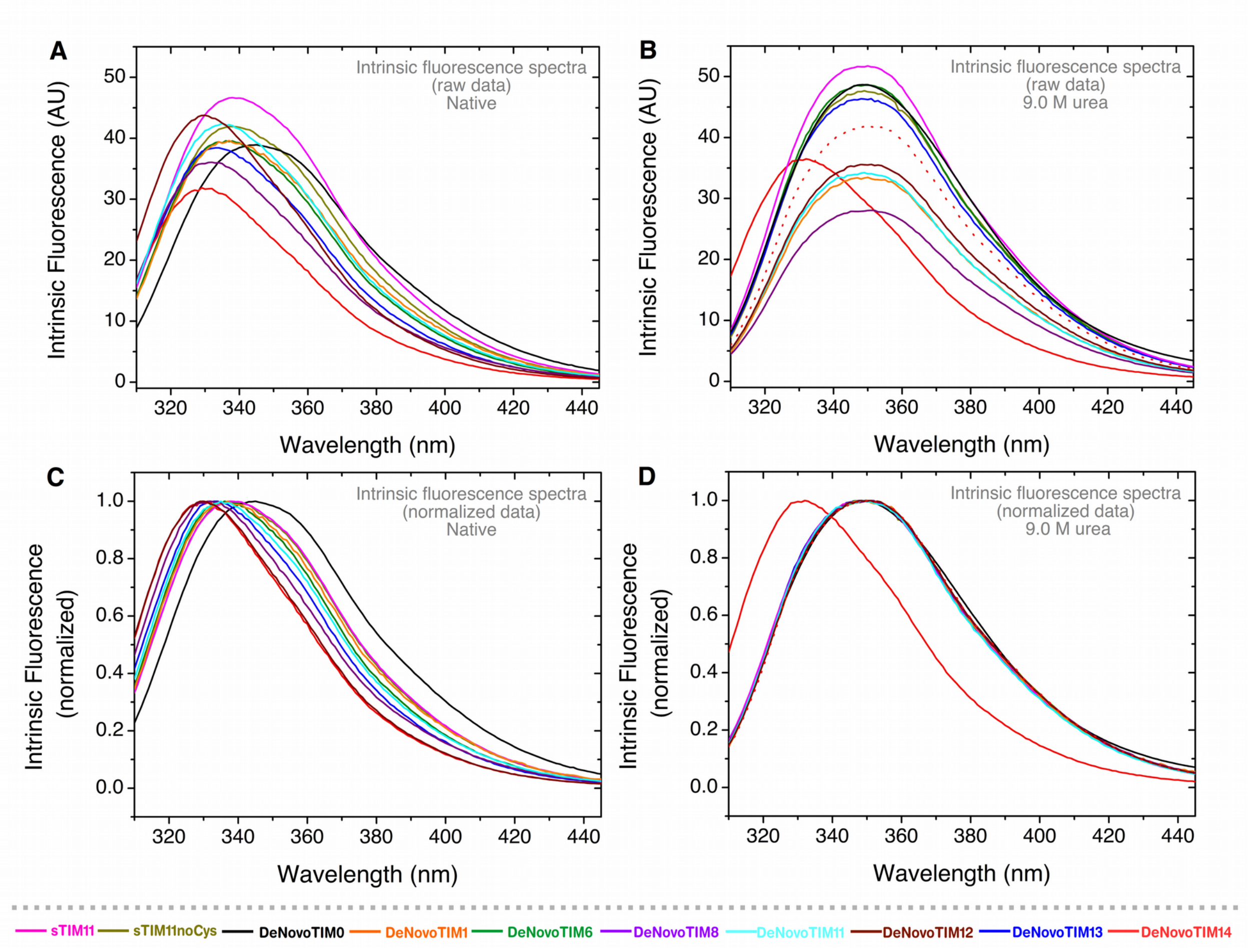


**Fig. S6. Intrinsic fluorescence spectra of DeNovoTIMs.** Native spectra are presented in panel A (raw data) and C (normalized data). Spectra obtained in 9.0 M urea are presented in panel B (raw data) and D (normalized data). Note that for DeNovoTIM14 in 9.0 M urea (solid red lines in panels B and D) the spectra are very similar to the native ones (solid red lines in panels A and C). In 7.0 M GdnHCl the protein is completely unfolded and its spectrum is identical to all other DeNovoTIMs unfolded in urea (dotted red line in panels C and D). In all experiments protein concentration was 0.4 mg $mL^{-1}$ in 10 mM sodium phosphate pH 8.

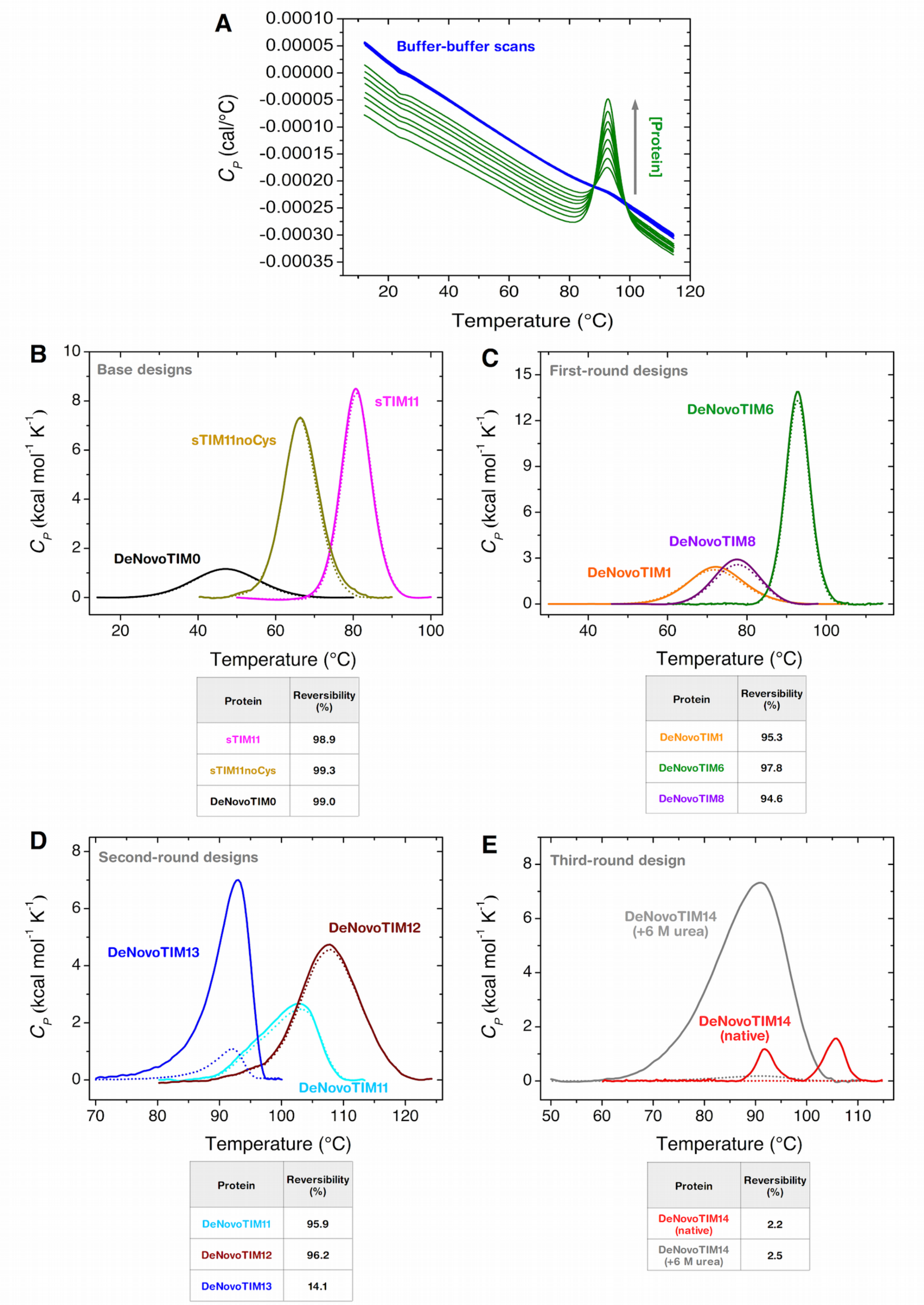


| Protein | Reversibility (%) |
|---|---|
| sTIM11 | 98.9 |
| sTIM11noCys | 99.3 |
| DeNovoTIM0 | 99.0 |

| Protein | Reversibility (%) |
|---|---|
| DeNovoTIM1 | 95.3 |
| DeNovoTIM6 | 97.8 |
| DeNovoTIM8 | 94.6 |

| Protein | Reversibility (%) |
|---|---|
| DeNovoTIM11 | 95.9 |
| DeNovoTIM12 | 96.2 |
| DeNovoTIM13 | 14.1 |

| Protein | Reversibility (%) |
|---|---|
| DeNovoTIM14 (native) | 2.2 |
| DeNovoTIM14 (+6 M urea) | 2.5 |

**Fig. S7. DSC instrument equilibration and thermal unfolding reversibility assessment of DeNovoTIMs. A)** Proper instrument equilibration was ascertained by performing two buffer-buffer scans before each protein-buffer scan. One example at different protein concentrations (DeNovoTIM6) is shown in the panel. **B-E)** Thermal unfolding reversibility was assessed with 1.0 mg protein mL$^{-1}$ and 60 K h$^{-1}$ in 10 mM sodium phosphate pH 8. Continuous lines show the first scan and dotted lines show the second scan collected after cooling down and reheating the sample. Reversibility % was calculated as the ratio of the calorimetric Δ*H* (area under the curve) recovered in the second scan and that obtained in the first scan ($\Delta H_{secondscan} / \Delta H_{firstscan}$) *100.

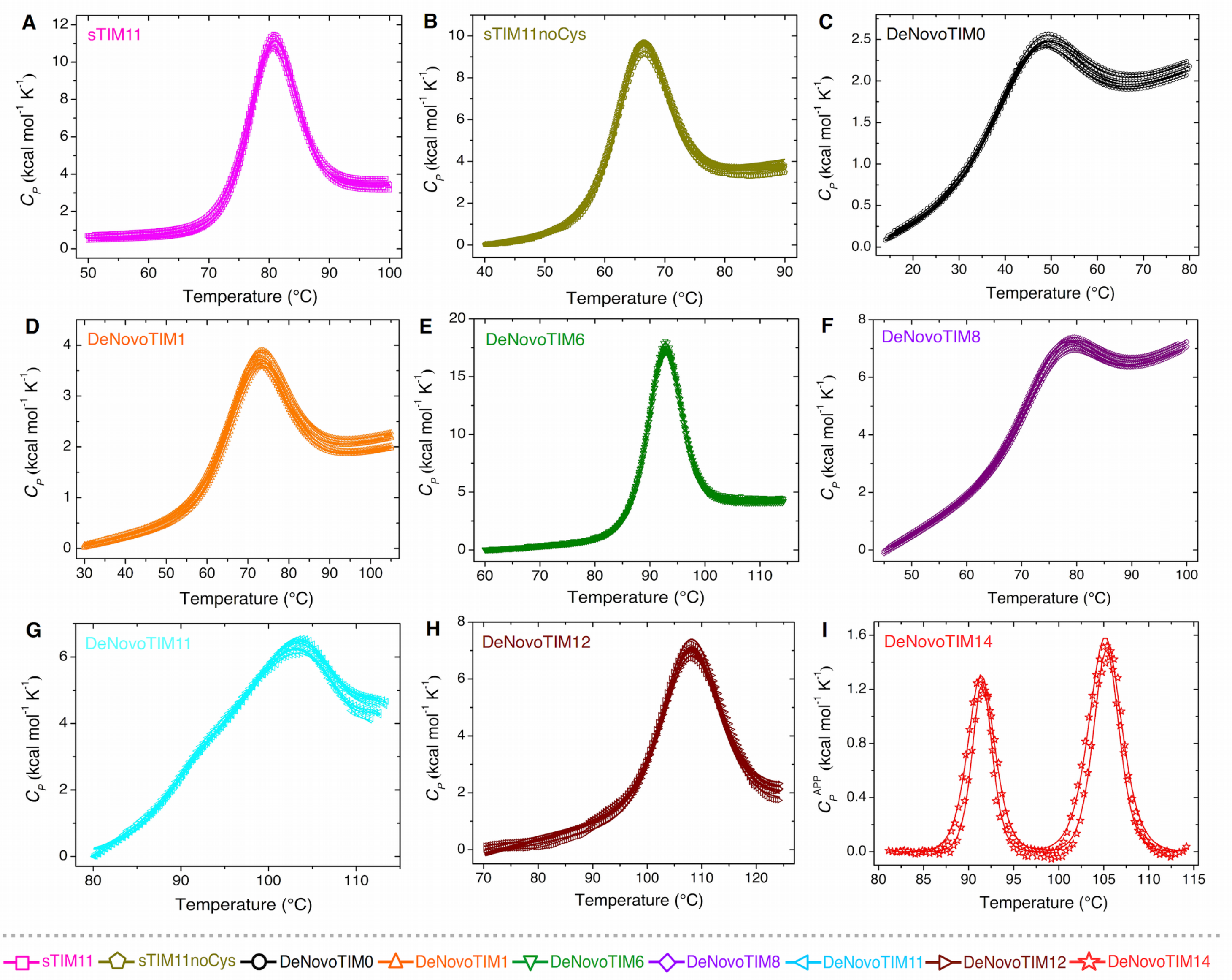


**Fig. S8. DSC endotherms of DeNovoTIMs**. Experiments were carried out at different protein concentrations (panels **A**-**H**: 0.25-2.5 mg mL$^{-1}$, panel **I**: 2.5 and 4.5 mg mL$^{-1}$). Open symbols show experimental data and solid lines are the best fits to a two-state model, except for DeNovoTIM14 where a non-two-state model with two transitions was used.

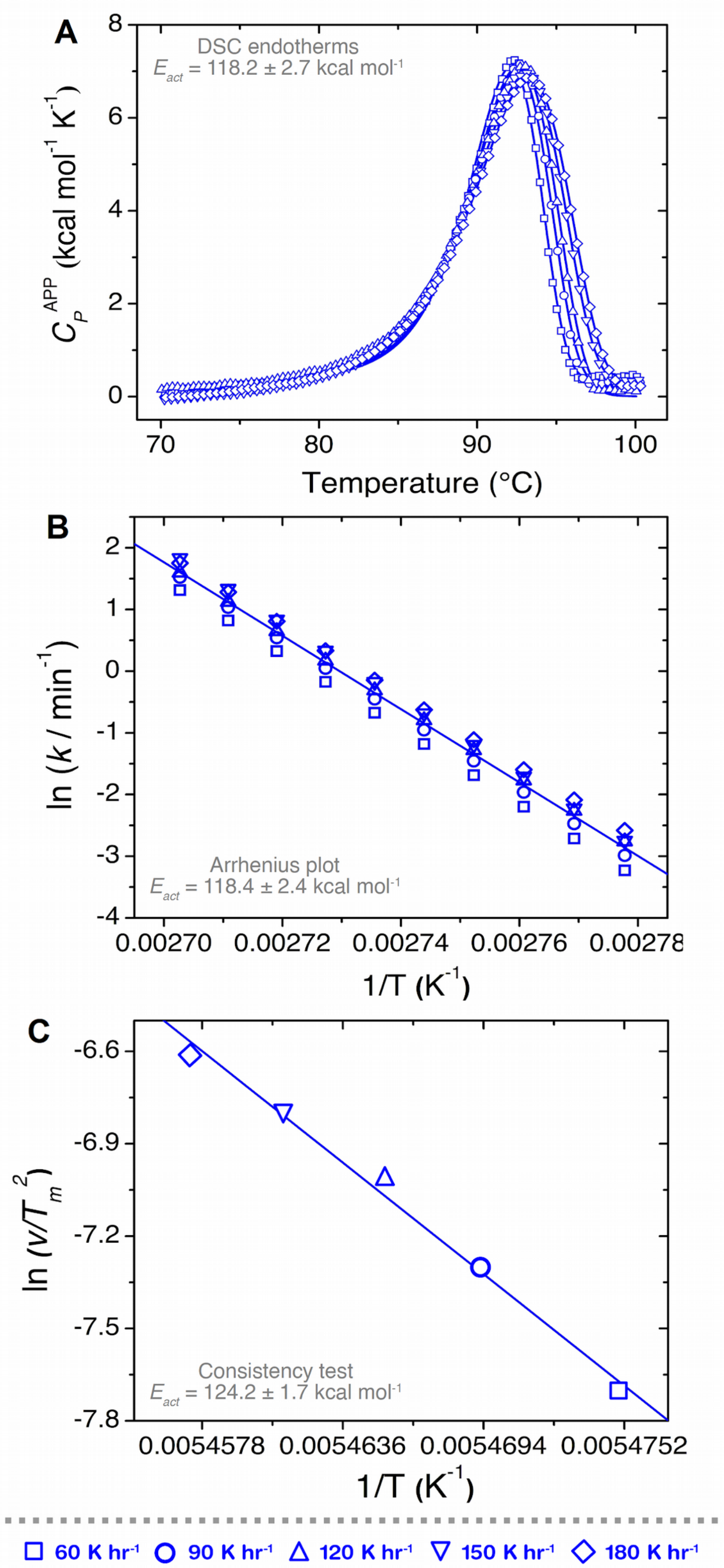


Fig. S9. Irreversible thermal unfolding of DeNovoTIM13. A) Endotherms at different scan rates (60 to 180 K $h^{-1}$). Lines represent the best fit to a two-state irreversible model. B) Arrhenius plot. The line shows the best fit to the Arrhenius equation ($R^2$: 0.98). C) Effect of the scan rate on $T_m$ ($R^2$: 0.99). In all experiments protein concentration was 0.5 mg $mL^{-1}$ in 10 mM sodium phosphate pH 8.

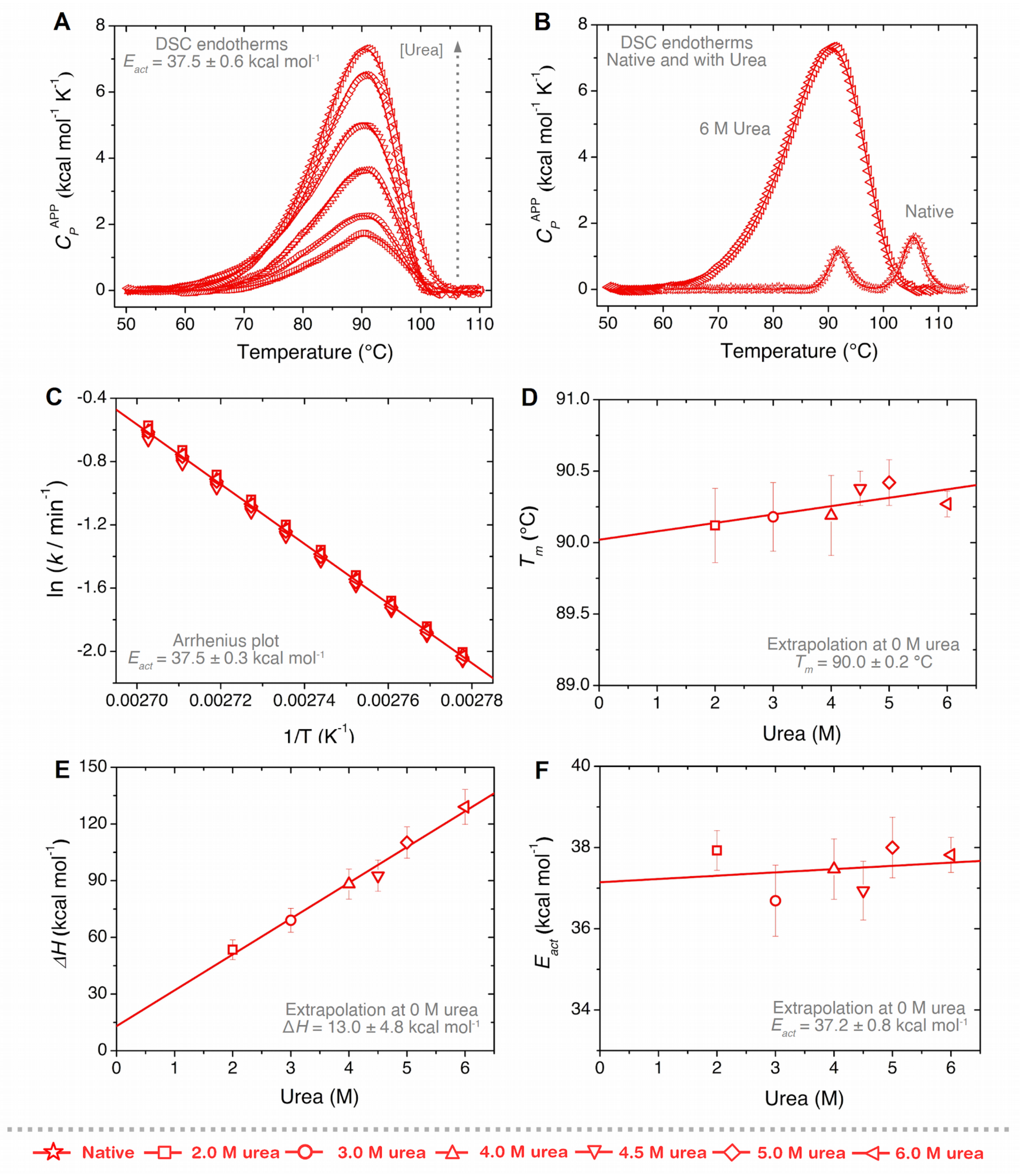


**Fig. S10. Irreversible thermal unfolding of DeNovoTIM14 in the presence of urea. A)** DSC endotherms at different urea concentrations (2.0 to 6.0 M). Lines represent the best fit to a two-state irreversible model. **B)** Comparison between DeNovoTIM14 endotherms in native conditions and at 6.0 M urea. **C)** Arrhenius plot. The line shows the best fit to the Arrhenius equation ($R^2$: 0.99). **D)** $T_m$ vs. urea concentration ($R^2$: 0.70). **E)** $\Delta H$ vs. urea concentration ($R^2$: 0.99). **F)** $E_{act}$ vs. urea concentration ($R^2$: 0.62). In all experiments protein concentration was 1 mg $mL^{-1}$ in 10 mM sodium phosphate pH 8.

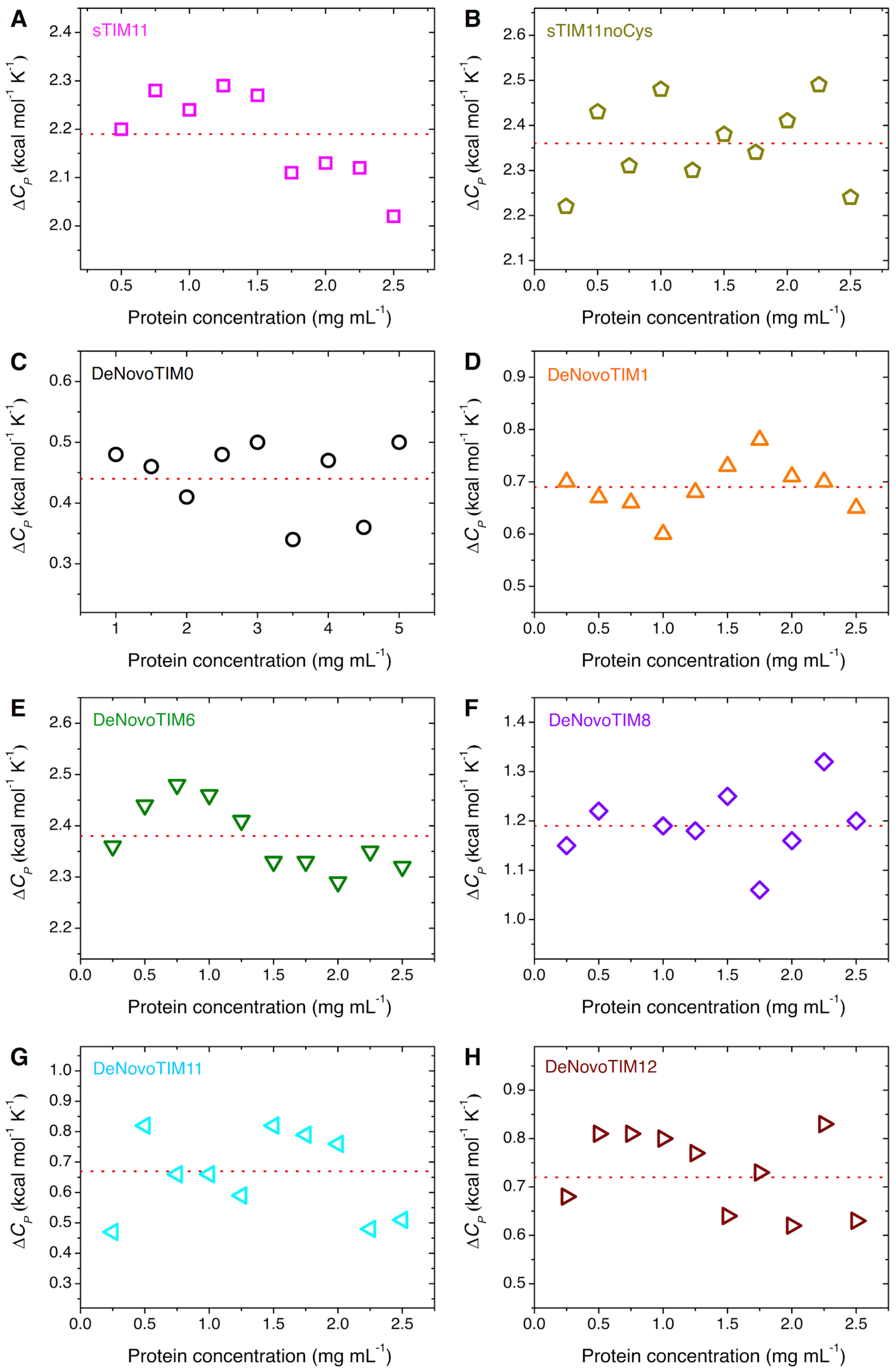


**Fig. S11. Δ$C_P$ determination of DeNovoTIMs at different protein concentrations**. DSC experiments (Fig. S8) were carried out in the range of 0.25-2.5 mg mL$^{-1}$ (panels **A**, **B**, **D**, **E**, **F**, **G**, **H**) or 1.0-5.0 mg mL$^{-1}$ (panel **C**: 1.0-5.0 mg mL$^{-1}$). Dotted red lines represent the average value for each protein.

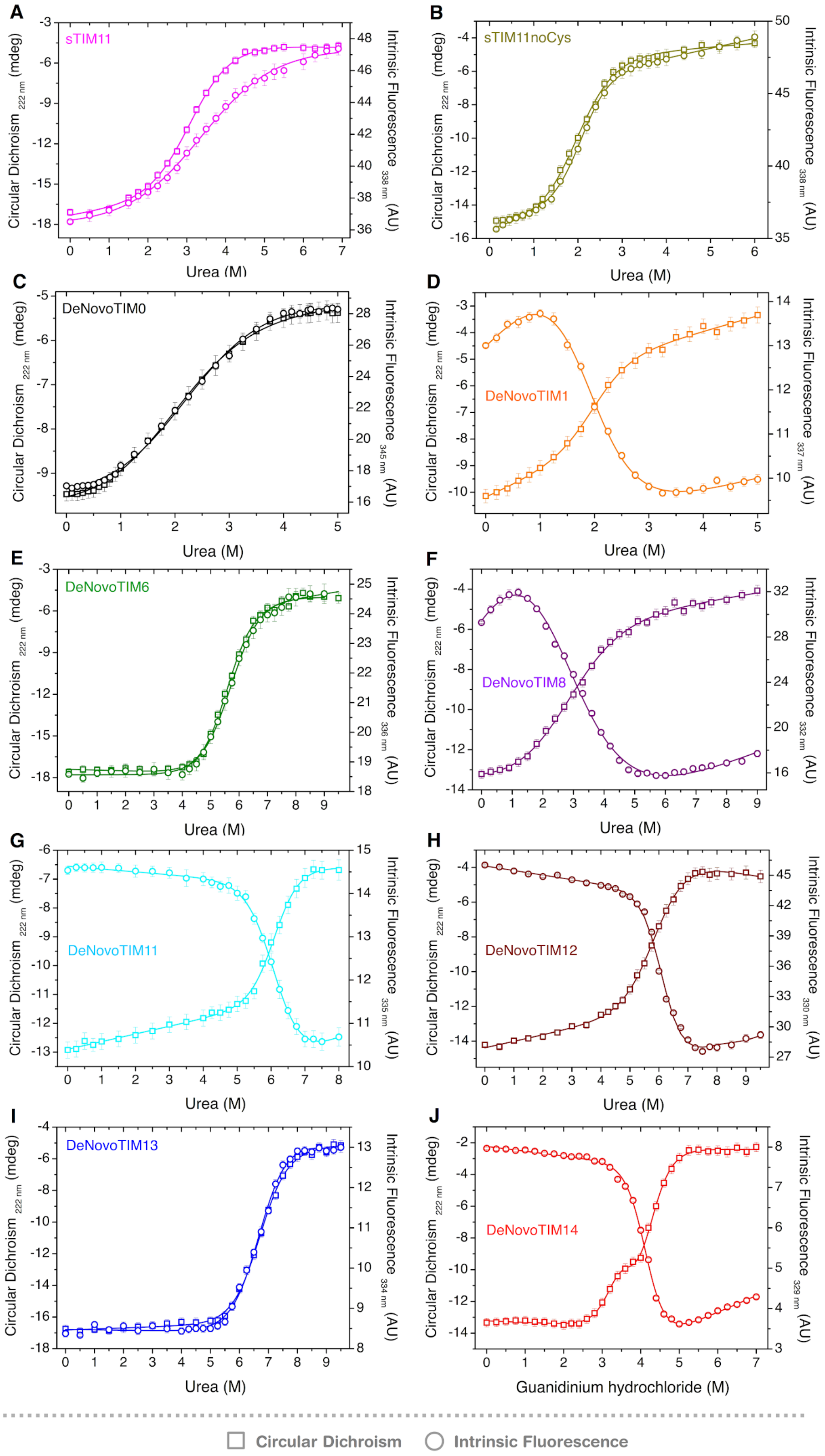


**Fig. S12. Chemical unfolding of DeNovoTIMs followed by CD and IF (raw data).** Lines are the best fits to a two-state model, except for DeNovoTIM14 where a three-state model was used. In all experiments protein concentration was 0.1 mg $mL^{-1}$ in 10 mM sodium phosphate pH 8.

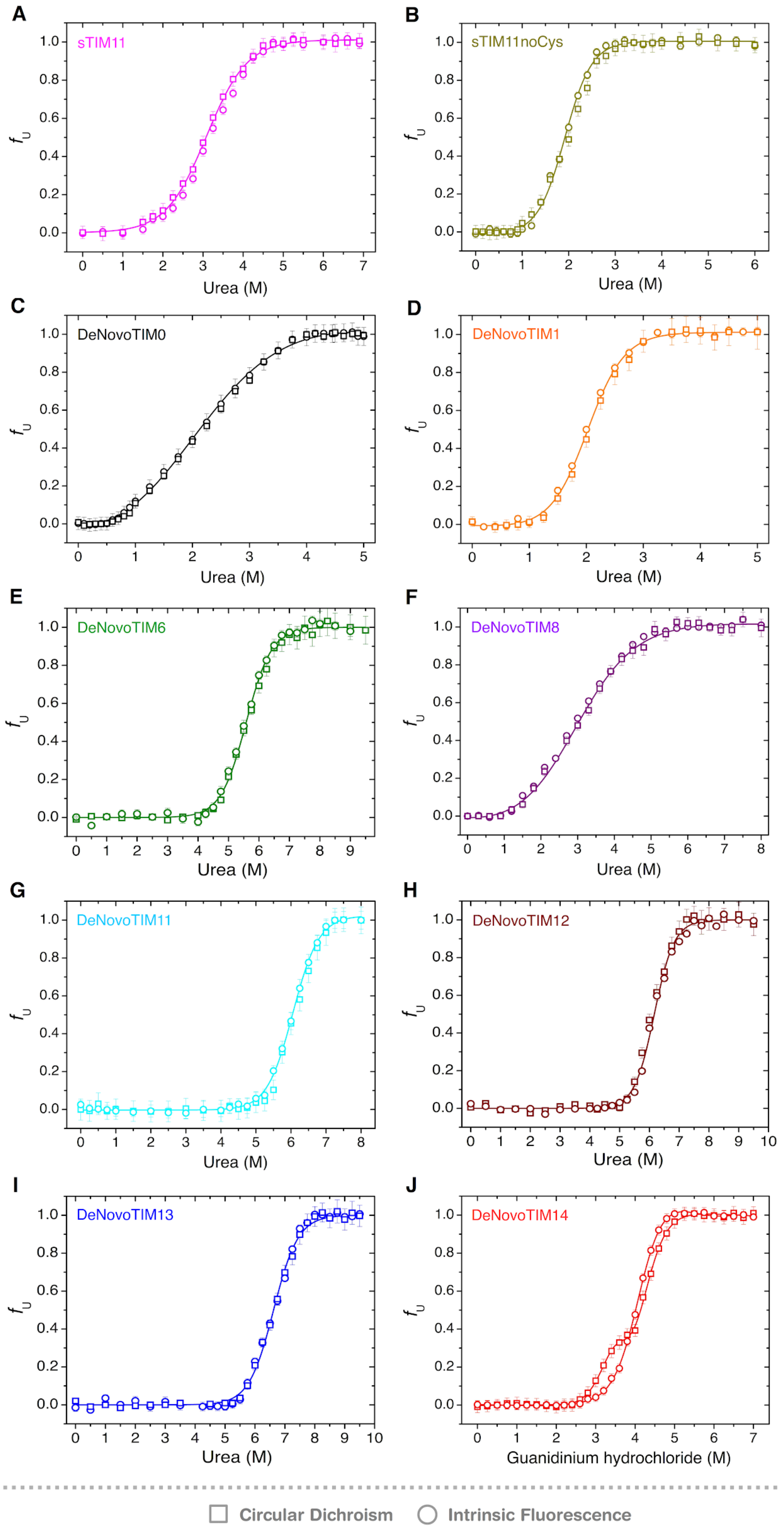


**Fig. S13. Chemical unfolding of DeNovoTIMs followed by CD and IF (normalized data).** Lines are the best fits to a two-state model, except for DeNovoTIM14 where a three-state model was used. In all experiments protein concentration was 0.1 mg mL$^{-1}$ in 10 mM sodium phosphate pH 8.

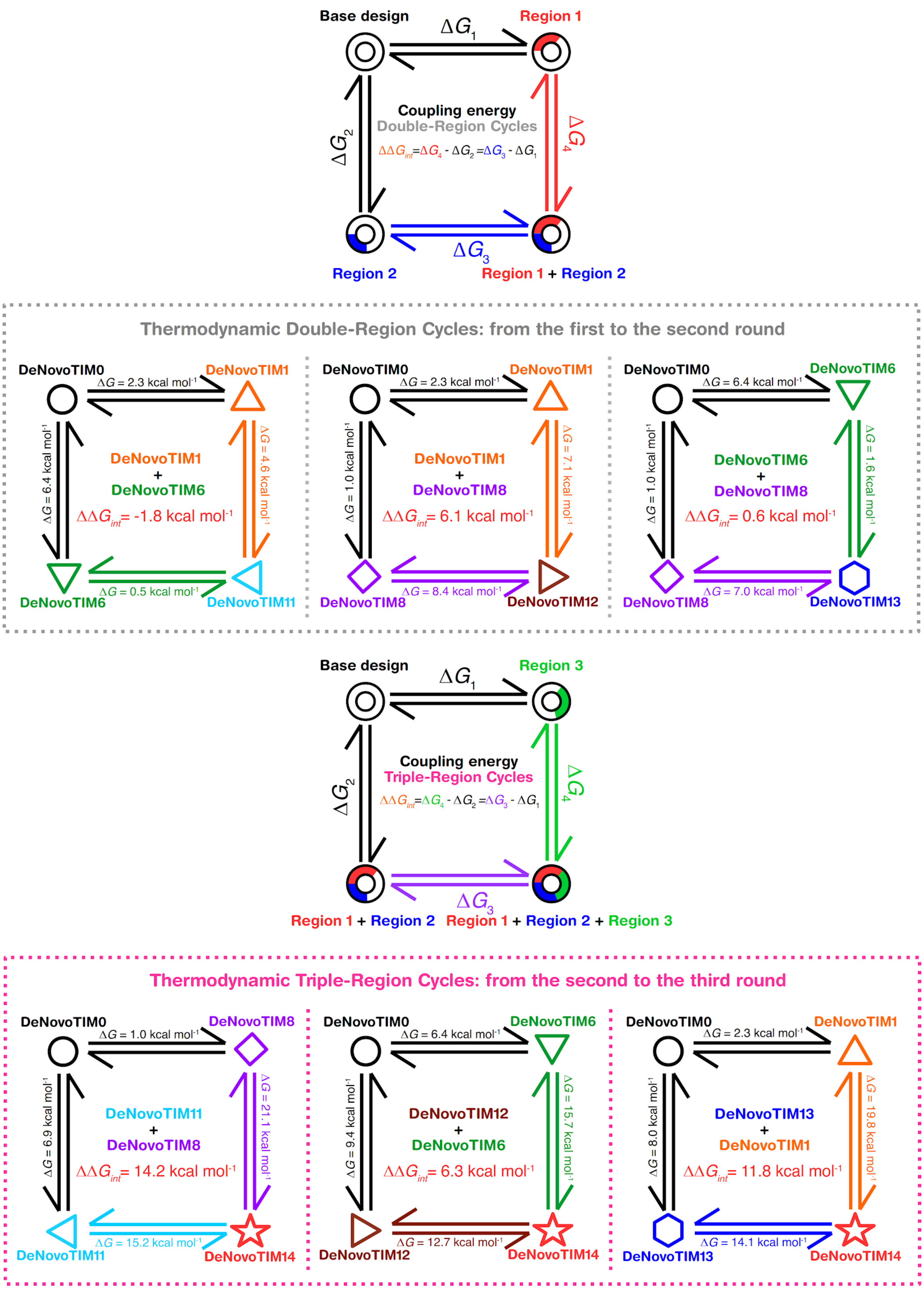


Fig. S14. Thermodynamic cycles for DeNovoTIMs. Thermodynamic cycles for double- and triple-region designs (top and bottom panels, respectively). Coupling energy ($\Delta\Delta G_{int}$) for each case is shown inside each cycle.

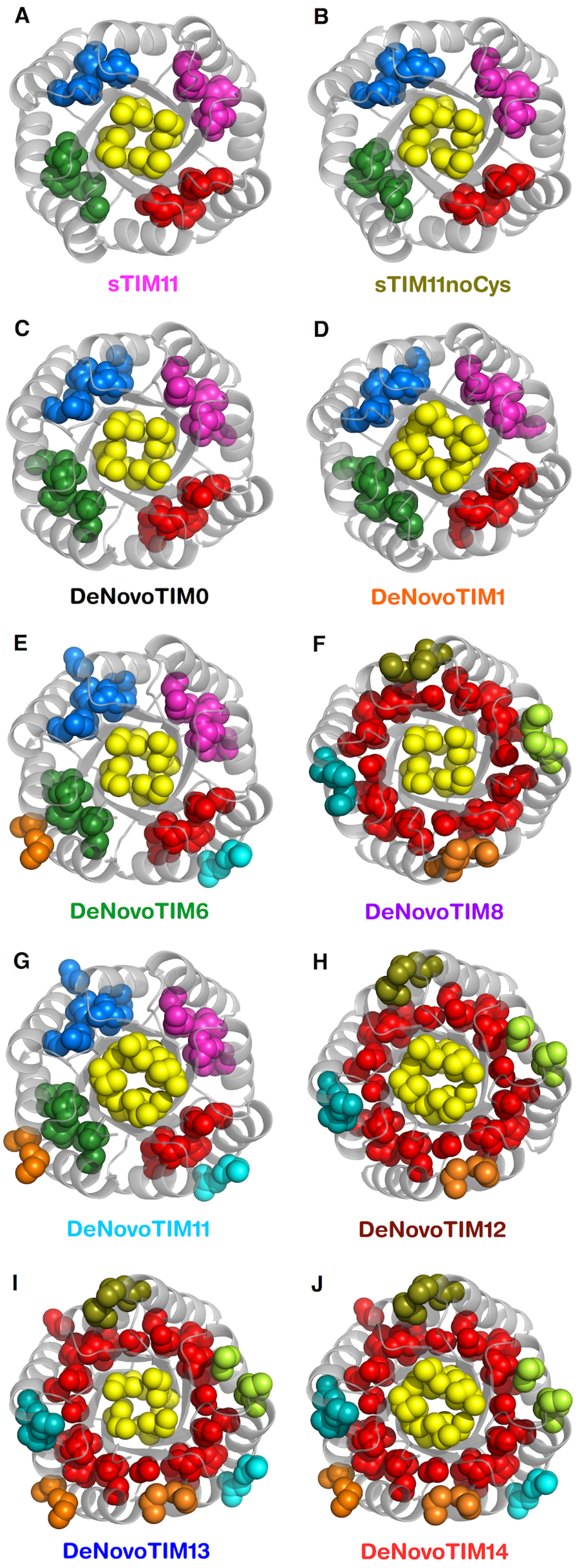


**Fig. S15. Hydrophobic clusters of DeNovoTIMs.** Clusters are shown in different colors. Hydrophobic clusters properties are reported in table S7.

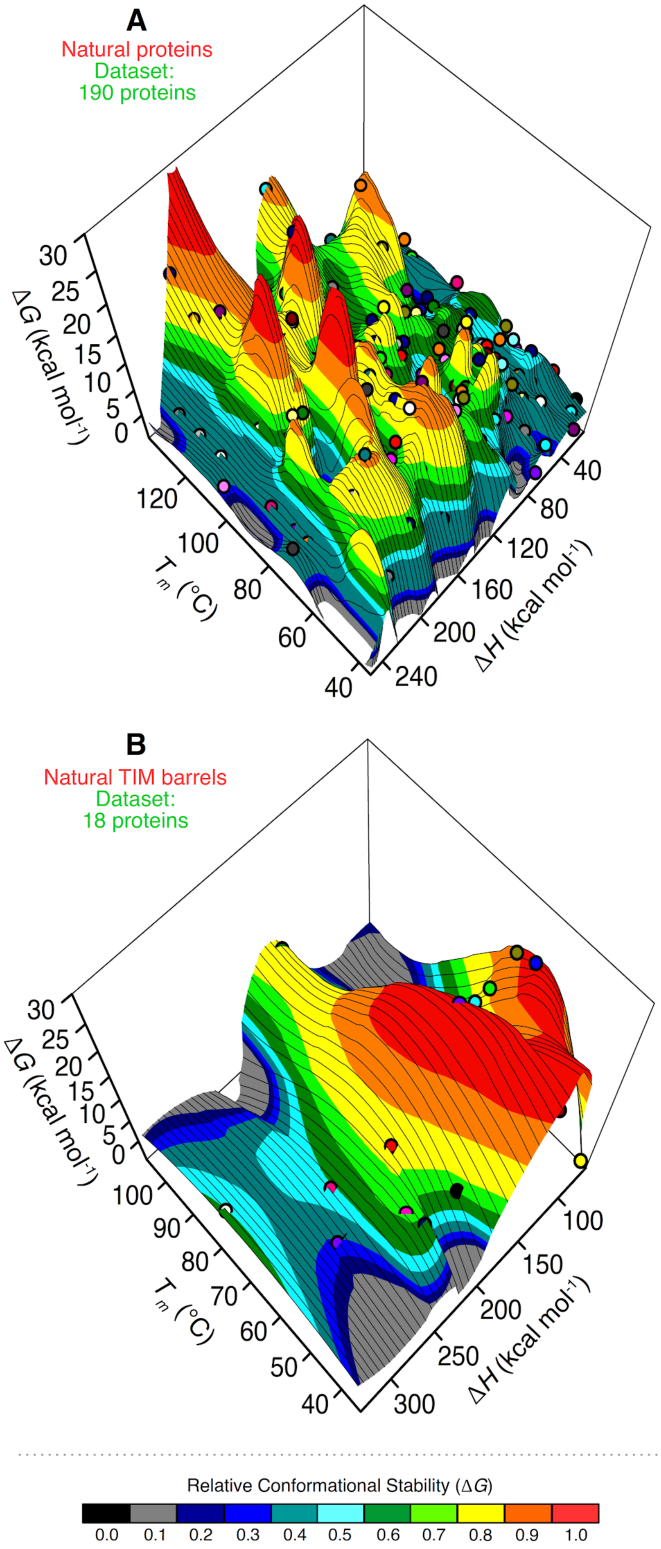


**Fig. S16. Stability landscape of natural proteins.** Panel A includes 190 natural proteins with different topologies varying in size from 42 to 572 residues (average: 173 residues). Panel B contains natural TIM barrels varying in size from 225 to 476 (average: 311 residues). The stability surface is colored according to normalized Δ*G* at 25 °C in 0.1 bins. Data for all non-redundant proteins presented here were obtained from ProThermDB database (*38, 39*).

### Supplementary tables

#### List of supplementary tables


- **Table S1:** Amino acid sequences of DeNovoTIM collection.
- **Table S2**: List of mutations in DeNovoTIM collection.
- **Table S3**: Aromatic residues in DeNovoTIM collection.
- **Table S4**: Sequence identity matrix of DeNovoTIM collection.
- **Table S5**: Biochemical and biophysical characterization of the DeNovoTIM collection.
- **Table S6**: Crystallographic data collection and refinement statistics for sTIM11noCys and DeNovoTIM structures.
- **Table S7**: Comparison of structural features between Rosetta models and three-dimensional structures.

Table S1. Amino acid sequences of DeNovoTIM collection.

| ***de novo*** **TIM barrel** | **Sequence** |
|---|---|
| sTIM11 | DKDEAWKCVEQLRREGATQIAYRSDDWRDLKEAWKKGADILIVDATDKDEAWKQVEQLRREGATQIAYRSDDWRDLKEAWKKGADILIVDAT<br>DKDEAWKQVEQLRREGATQIAYRSDDWRDLKEAWKKGADILIVDATDKDEAWKQVEQLRREGATQIAYRSDDWRDLKEAWKKGADILICDAT |
| sTIM11noCys | DKDEAWKQVEQLRREGATQIAYRSDDWRDLKEAWKKGADILIVDATDKDEAWKQVEQLRREGATQIAYRSDDWRDLKEAWKKGADILIVDAT<br>DKDEAWKQVEQLRREGATQIAYRSDDWRDLKEAWKKGADILIVDATDKDEAWKQVEQLRREGATQIAYRSDDWRDLKEAWKKGADILIVDAT |
| DeNovoTIM0 | DKDEAWKQVEQLRREGATQIAYRSDDWRDLKEAVKKGGDILIVDATDKDEAWKQVEQLRREGATQIAYRSDDWRDLKEAVKKGGDILIVDAT<br>DKDEAWKQVEQLRREGATQIAYRSDDWRDLKEAVKKGGDILIVDATDKDEAWKQVEQLRREGATQIAYRSDDWRDLKEAVKKGGDILIVDAT |
| DeNovoTIM1 | DKDEAWKQVEQLRREGATQIVYISDDWRDLKEAVKKGGDILIVDATDKDEAWKQVEQLRREGATQIWYISDDWRDLKEAVKKGGDVLIVDAT<br>DKDEAWKQVEQLRREGATQIVYISDDWRDLKEAVKKGGDILIVDATDKDEAWKQVEQLRREGATQIWYISDDWRDLKEAVKKGGDVLIVDAT |
| DeNovoTIM2 | DKDEAWKQVEQLRREGATQIVYQSDDWRDLKEAVKKGGDILMVDATDKDEAWKQVEQLRREGATQIMYISDDWRDLKEAVKKGGDFLVVDAT<br>DKDEAWKQVEQLRREGATQIVYQSDDWRDLKEAVKKGGDILMVDATDKDEAWKQVEQLRREGATQIMYISDDWRDLKEAVKKGGDFLVVDAT |
| DeNovoTIM3 | DKDEAWKQVEQLRREGATQIMYVSDDWRDLKEAVKKGGDSLLVDATDKDEAWKQVEQLRREGATQILYKSDDWRDLKEAVKKGGDTLVVDAT<br>DKDEAWKQVEQLRREGATQIMYVSDDWRDLKEAVKKGGDSLLVDATDKDEAWKQVEQLRREGATQILYKSDDWRDLKEAVKKGGDTLVVDAT |
| DeNovoTIM4 | DKDEAWKQVEQLRREGATQIMYFSDDWRDLKEAVKKGGDLLVVDATDKDEAWKQVEQLRREGATQILYISDDWRDLKEAVKKGGDILLVDAT<br>DKDEAWKQVEQLRREGATQIMYFSDDWRDLKEAVKKGGDLLVVDATDKDEAWKQVEQLRREGATQILYISDDWRDLKEAVKKGGDILLVDAT |
| DeNovoTIM5 | DKDEAWKQVEQLRRLGATQIAYRSDDWRDLREAVKKGGDILIVDATDKDEAWKQVEQLRRLGATQIAYRSDDWRDLREAVKKGGDILIVDAT<br>DKDEAWKQVEQLRRLGATQIAYRSDDWRDLREAVKKGGDILIVDATDKDEAWKQVEQLRRLGATQIAYRSDDWRDLREAVKKGGDILIVDAT |
| DeNovoTIM6 | DKDEAWKQVEILRRLGAKQIAYRSDDWRDLQEALKKGGDILIVDATDKDEAWKQVEILRRLGAKQIAYRSDDWRDLQEALKKGGDILIVDAT<br>DKDEAWKQVEILRRLGAKQIAYRSDDWRDLQEALKKGGDILIVDATDKDEAWKQVEILRRLGAKQIAYRSDDWRDLQEALKKGGDILIVDAT |
| DeNovoTIM7 | DKDEAWKQVEILRRLGAKQIAYRSDDWRDLDEARKKGGDILIVDATDKDEAWKQVEILRRLGAKQIAYRSDDWRDLDEARKKGGDILIVDAT<br>DKDEAWKQVEILRRLGAKQIAYRSDDWRDLDEARKKGGDILIVDATDKDEAWKQVEILRRLGAKQIAYRSDDWRDLDEARKKGGDILIVDAT |
| DeNovoTIM8 | DVDEMLKQVEQLRREGATQIAVRSDDWRILKEAVKKGGDILIVDATDVDEMLKQVEQLRREGATQIAVRSDDWRILKEAVKKGGDILIVDAT<br>DVDEMLKQVEQLRREGATQIAVRSDDWRILKEAVKKGGDILIVDATDVDEMLKQVEQLRREGATQIAVRSDDWRILKEAVKKGGDILIVDAT |
| DeNovoTIM9 | DPDEAQKQVEQLRREGATQIAIRSDDWRYLKEAVKKGGDILIVDATDPDEAQKQVEQLRREGATQIAIRSDDWRYLKEAVKKGGDILIVDAT<br>DPDEAQKQVEQLRREGATQIAIRSDDWRYLKEAVKKGGDILIVDATDPDEAQKQVEQLRREGATQIAIRSDDWRYLKEAVKKGGDILIVDAT |
| DeNovoTIM10 | DSDEAIKQVEQLRREGATQIAVRMDDWRKLKEAVKKGGDILIVDATDSDEAIKQVEQLRREGATQIAVRMDDWRKLKEAVKKGGDILIVDAT<br>DSDEAIKQVEQLRREGATQIAVRMDDWRKLKEAVKKGGDILIVDATDSDEAIKQVEQLRREGATQIAVRMDDWRKLKEAVKKGGDILIVDAT |
| DeNovoTIM11 | DKDEAWKQVEILRRLGAKQIVYISDDWRDLQEALKKGGDILIVDATDKDEAWKQVEILRRLGAKQIWYISDDWRDLQEALKKGGDVLIVDAT<br>DKDEAWKQVEILRRLGAKQIVYISDDWRDLQEALKKGGDILIVDATDKDEAWKQVEILRRLGAKQIWYISDDWRDLQEALKKGGDVLIVDAT |
| DeNovoTIM12 | DVDEMLKQVEQLRREGATQIVVISDDWRILKEAVKKGGDILIVDATDVDEMLKQVEQLRREGATQIWVISDDWRILKEAVKKGGDVLIVDAT<br>DVDEMLKQVEQLRREGATQIVVISDDWRILKEAVKKGGDILIVDATDVDEMLKQVEQLRREGATQIWVISDDWRILKEAVKKGGDVLIVDAT |
| DeNovoTIM13 | DVDEMLKQVEILRRLGAKQIAVRSDDWRILQEALKKGGDILIVDATDVDEMLKQVEILRRLGAKQIAVRSDDWRILQEALKKGGDILIVDAT<br>DVDEMLKQVEILRRLGAKQIAVRSDDWRILQEALKKGGDILIVDATDVDEMLKQVEILRRLGAKQIAVRSDDWRILQEALKKGGDILIVDAT |
| DeNovoTIM14 | DVDEMLKQVEILRRLGAKQIVVISDDWRILQEALKKGGDILIVDATDVDEMLKQVEILRRLGAKQIWVISDDWRILQEALKKGGDVLIVDAT<br>DVDEMLKQVEILRRLGAKQIVVISDDWRILQEALKKGGDILIVDATDVDEMLKQVEILRRLGAKQIWVISDDWRILQEALKKGGDVLIVDAT |

Table S2. List of mutations in DeNovoTIM collection.

| *de novo* TIM barrel | Symmetry (fold) | Number of mutations in each modular unit[a] | Total number of mutations in the Sequence | Mutations added in each modular unit[a] |
|---|---|---|---|---|
| sTIM11 | pseudo 4 | ----- | ----- | ----- |
| sTIM11noCys | 4 | 2 (non-symmetric) | 2 | C8Q, C181V |
| DeNovoTIM0 | 4 | 2 | 8 | W34V, A38G |
| DeNovoTIM1 | 2 | 5 | 10 | A21V, R23I, A67W, R69I, I86V |
| DeNovoTIM2 | 2 | 7 | 14 | A21V, R23Q, I42M, A67M, R69I, I86F, I88V |
| DeNovoTIM3 | 2 | 8 | 16 | A21M, R23V, I40S, I42L, A67L, R69K, I86T, I88V |
| DeNovoTIM4 | 2 | 7 | 14 | A21M, R23F, I40L, I42V, A67L, R69I, I88L |
| DeNovoTIM5 | 4 | 2 | 8 | E15L, K31R |
| DeNovoTIM6 | 4 | 5 | 20 | Q11I, E15L, T18K, K31Q, V34L |
| DeNovoTIM7 | 4 | 5 | 20 | Q11I, E15L, T18K, K31D, V34R |
| DeNovoTIM8 | 4 | 5 | 20 | K2V, A5M, W6L, Y22V, D29I |
| DeNovoTIM9 | 4 | 4 | 16 | K2P, W6Q, Y22I, D29Y |
| DeNovoTIM10 | 4 | 5 | 20 | K2S, W6I, Y22V, S24M, D29K |
| DeNovoTIM11 | 2 | 15 | 30 | Q11I, E15L, T18K, A21V, R23I, K31Q, V34L, Q57I, E61L, A67W, T64K, R69I, K77Q, V80L, I86V |
| DeNovoTIM12 | 2 | 15 | 30 | K2V, A5M, W6L, A21V, Y22V, R23I, D29I, K48V, A51M, W52L, A67W, Y68V, R69I, D75I, I86V |
| DeNovoTIM13 | 4 | 10 | 40 | K2V, A5M, W6L, Q11I, E15L, T18K, Y22V, D29I, K31Q, V34L |
| DeNovoTIM14 | 2 | 25 | 50 | K2V, A5M, W6L, Q11I, E15L, T18K, A21V, Y22V, R23I, D29I, K31Q, V34L, K48V, A51M, W52L, Q57I, E61L, T64K, A67W, Y68V, R69I, D75I, K77Q, V80L, I86V |

[a] Only the mutations incorporated in each design are shown. See Fig. S2 for the complete sequence alignment.

Table S3. Aromatic residues in DeNovoTIM collection.

| *de novo* TIM barrel | Number of Tyr residues in each modular unit | Total number of Trp residues | Position of Trp residues in the modular unit | Number of Tyr residues in each modular unit | Total number of Tyr residues | Position of Tyr residues in the modular unit |
|---|---|---|---|---|---|---|
| sTIM11 | 3 | 12 | W6, W27, W34 | 1 | 4 | Y22 |
| sTIM11noCys | 3 | 12 | W6, W27, W34 | 1 | 4 | Y22 |
| DeNovoTIM0 | 2 | 8 | W6, W27 | 1 | 4 | Y22 |
| DeNovoTIM1 | 5 | 10 | W6, W27, W52, W67, W73 | 2 | 4 | Y22, Y68 |
| DeNovoTIM2 | 4 | 8 | W6, W27, W52, W73 | 2 | 4 | Y22, Y68 |
| DeNovoTIM3 | 4 | 8 | W6, W27, W52, W74 | 2 | 4 | Y22, Y68 |
| DeNovoTIM4 | 4 | 8 | W6, W27, W52, W75 | 2 | 4 | Y22, Y68 |
| DeNovoTIM5 | 2 | 8 | W6, W27 | 1 | 4 | Y22 |
| DeNovoTIM6 | 2 | 8 | W6, W27 | 1 | 4 | Y22 |
| DeNovoTIM7 | 2 | 8 | W6, W27 | 1 | 4 | Y22 |
| DeNovoTIM8 | 1 | 4 | W27 | 0 | 0 | ----- |
| DeNovoTIM9 | 1 | 4 | W27 | 0 | 0 | ----- |
| DeNovoTIM10 | 1 | 4 | W27 | 0 | 0 | ----- |
| DeNovoTIM11 | 5 | 10 | W6, W27, W52, W67, W73 | 2 | 4 | Y22, Y68 |
| DeNovoTIM12 | 5 | 10 | W6, W27, W52, W67, W73 | 0 | 0 | ----- |
| DeNovoTIM13 | 2 | 8 | W6, W27 | 0 | 0 | ----- |
| DeNovoTIM14 | 5 | 10 | W6, W27, W52, W67, W73 | 0 | 0 | ----- |

Table S4. Sequence identity matrix of DeNovoTIM collection.

| | sTIM11 | sTIM11noCys | DeNovoTIM0 | DeNovoTIM1 | DeNovoTIM2 | DeNovoTIM3 | DeNovoTIM4 | DeNovoTIM5 | DeNovoTIM6 | DeNovoTIM7 | DeNovoTIM8 | DeNovoTIM9 | DeNovoTIM10 | DeNovoTIM11 | DeNovoTIM12 | DeNovoTIM13 | DeNovoTIM14 |
|---|---|---|---|---|---|---|---|---|---|---|---|---|---|---|---|---|---|
| sTIM11 | | | | | | | | | | | | | | | | | |
| sTIM11noCys | 99 | | | | | | | | | | | | | | | | |
| DeNovoTIM0 | 95 | 96 | | | | | | | | | | | | | | | |
| DeNovoTIM1 | 89 | 90 | 95 | | | | | | | | | | | | | | |
| DeNovoTIM2 | 87 | 88 | 92 | 95 | | | | | | | | | | | | | |
| DeNovoTIM3 | 86 | 87 | 91 | 91 | 92 | | | | | | | | | | | | |
| DeNovoTIM4 | 87 | 88 | 92 | 92 | 92 | 94 | | | | | | | | | | | |
| DeNovoTIM5 | 90 | 91 | 96 | 90 | 88 | 87 | 88 | | | | | | | | | | |
| DeNovoTIM6 | 86 | 87 | 89 | 84 | 82 | 80 | 82 | 91 | | | | | | | | | |
| DeNovoTIM7 | 86 | 87 | 89 | 84 | 82 | 80 | 82 | 91 | 96 | | | | | | | | |
| DeNovoTIM8 | 84 | 85 | 89 | 84 | 82 | 80 | 82 | 85 | 78 | 78 | | | | | | | |
| DeNovoTIM9 | 86 | 87 | 91 | 86 | 84 | 83 | 84 | 87 | 80 | 80 | 89 | | | | | | |
| DeNovoTIM10 | 84 | 85 | 89 | 84 | 82 | 80 | 82 | 85 | 78 | 78 | 89 | 89 | | | | | |
| DeNovoTIM11 | 80 | 82 | 84 | 89 | 84 | 80 | 82 | 86 | 95 | 90 | 73 | 75 | 73 | | | | |
| DeNovoTIM12 | 78 | 79 | 84 | 89 | 84 | 80 | 82 | 79 | 73 | 73 | 95 | 84 | 84 | 78 | | | |
| DeNovoTIM13 | 75 | 76 | 78 | 73 | 71 | 70 | 71 | 80 | 89 | 85 | 89 | 78 | 78 | 84 | 84 | | |
| DeNovoTIM14 | 70 | 71 | 73 | 78 | 73 | 70 | 71 | 75 | 84 | 79 | 84 | 73 | 73 | 89 | 89 | 95 | |

Table S5. Biochemical and biophysical characterization of DeNovoTIM collection.

| *de novo* TIM barrel | Expression/Solubility properties | | Spectroscopic properties: Circular Dichroism – Predicted secondary structure content (%)[c] | | | Spectroscopic properties: Intrinsic Fluorescence[d] | | | | | | Hydrodynamic properties[e] | | | | |
|---|---|---|---|---|---|---|---|---|---|---|---|---|---|---|---|---|
| | Soluble overexpression[a] | Aggregation-prone[b] | α-helix | β-strand | Random coil | Native $\lambda_{max}$ (nm) | Unfolded $\lambda_{max}$ (nm) | $\Delta\lambda_{max\ N\text{-}U}$ (nm) | Native SCM (nm) | Unfolded SCM (nm) | $\Delta SCM_{N\text{-}U}$ (nm) | Theoretical MW (Da) | Experimental MW (Da) | Exp./ Theor. ratio | Oligomeric state | Stokes radius (Å) |
| sTIM11 | +++ | + | 46.3 (46.0) | 19.8 (20.0) | 34.0 (34.0) | 338 | 350 | 12 | 354 | 360 | 7 | 22854 | 22017 ± 465 | 1.0 | Monomer | 23.2 ± 0.5 |
| sTIM11noCys | +++ | + | 47.6 (48.1) | 19.2 (22.8) | 33.2 (29.1) | 338 | 349 | 11 | 353 | 361 | 7 | 22875 | 20670 ± 146 | 0.9 | Monomer | 22.6 ± 0.1 |
| DeNovoTIM0 | ++ | + | 41.4 | 30.7 | 27.9 | 345 | 352 | 7 | 357 | 362 | 5 | 22471 | 29827 ± 309 | 1.3 | Monomer | 26.1 ± 0.3 |
| DeNovoTIM1 | ++ | ++ | 42.7 | 29.3 | 27.9 | 337 | 350 | 13 | 354 | 360 | 6 | 22557 | 21017 ± 637 | 0.9 | Monomer | 22.8 ± 0.8 |
| DeNovoTIM6 | +++ | + | 46.5 (43.4) | 18.8 (20.7) | 34.8 (35.9) | 336 | 350 | 14 | 352 | 361 | 9 | 22568 | 22000 ± 440 | 1.0 | Monomer | 23.2 ± 0.5 |
| DeNovoTIM8 | ++ | ++ | 47.1 | 22.3 | 30.6 | 332 | 351 | 19 | 348 | 361 | 13 | 22039 | 20518 ± 360 | 0.9 | Monomer | 22.6 ± 0.3 |
| DeNovoTIM11 | + | +++ | 46.9 | 22.9 | 30.0 | 335 | 349 | 14 | 353 | 362 | 9 | 22598 | 20920 ± 572 | 0.9 | Monomer | 22.3 ± 0.6 |
| DeNovoTIM12 | +++ | + | 45.5 | 24.7 | 29.7 | 330 | 351 | 21 | 347 | 360 | 13 | 22125 | 20029 ± 760 | 0.9 | Monomer | 23.1 ± 0.9 |
| DeNovoTIM13 | +++ | +++ | 48.3 (48.8) | 18.1 (21.8) | 33.5 (29.4) | 334 | 351 | 17 | 351 | 360 | 8 | 22080 | 21601 ± 725 | 1.0 | Monomer | 22.7 ± 0.7 |
| DeNovoTIM14 | + | +++ | 50.3 | 18.7 | 31.1 | 329 | 350 (331) | 21 | 346 | 362 (349) | 15 | 22166 | 25357 ± 1833 | 1.2 | Monomer | 24.7 ± 1.8 |

[a] Symbols represent the amount of protein purified from the soluble fraction: +++ >10 mg $L^{-1}$ culture, ++ 5-10 mg $L^{-1}$ culture, + 2-5 mg $L^{-1}$ culture.

[b] Symbols represent the protein propensity to aggregate: +++ aggregation visible within the first week after purification, ++ aggregation visible between 1-2 weeks after purification, + not visible aggregation after at least 1 month after purification.

[c] numbers within brackets are the calculated value from the three-dimensional structure.

[d] unfolded state represents the spectra at 9.0 M urea.

[e] ± indicate the standard deviation calculated from 4 different experiments at protein concentrations of 0.01, 0.5, 1.0, and 2.0 mg $mL^{-1}$.

Table S6. Crystallographic data collection and refinement statistics for sTIM11noCys and DeNovoTIM structures.

| *de novo* **TIM barrel** | **sTIM11noCys** | **DeNovoTIM6** | **DeNovoTIM13** |
|---|---|---|---|
| **Data collection[a]** | | | |
| PDB ID | 6YQY | 6Z2I | 6YQX |
| Crystallization condition | 0.2 M Ammonium sulfate<br>0.1 M Trisodium citrate pH: 5.6<br>25% w/v PEG 4000 | 0.095 M Sodium citrate pH: 5.0<br>19% v/v Isopropanol<br>25% w/v PEG 4000<br>5% v/v Glycerol | 0.17 M Sodium acetate<br>0.085 M TRIS pH: 8.5<br>25.5% w/v PEG 4000<br>15% v/v Glycerol |
| Resolution range (Å) | 47.15 – 1.88 (1.94 – 1.88) | 30.93 – 2.90 (3.01 – 2.90) | 44.79 – 1.64 (1.70 – 1.64) |
| Space group | $P\ 4_1\ 2_1\ 2$ (92) | P 4 (75) | $P\ 3_2$ (145) |
| Unit cell dimensions | | | |
| a, b, c, (Å) | 50.45, 50.45, 132.41 | 87.48, 87.48, 44.26 | 51.72, 51.72, 63.97 |
| α, β, γ (°) | 90, 90, 90 | 90, 90, 90 | 90, 90, 120 |
| Total reflections | 342791 (32681) | 56016 (3033) | 79664 (7327) |
| Unique reflections | 14703 (1410) | 7620 (389) | 23407 (2280) |
| Multiplicity | 23.3 (23.2) | 7.4 (7.8) | 3.4 (3.2) |
| Completeness (%) | 99.9 (99.0) | 99.8 (100.0) | 99.5 (97.6) |
| Mean I / sigma (I) | 27.0 (1.9) | 4.3 (1.0) | 14.3 (1.5) |
| R-merge | 0.061 (1.281) | 0.180 (1.195) | 0.042 (0.552) |
| CC1/2 | 1.000 (0.810) | 0.940 (0.773) | 0.998 (0.659) |
| Average mosaicity (deg) | 0.109 | 0.179 | 0.116 |
| **Refinement statistics[a]** | | | |
| $R_{work}$ , $R_{free}$ | 0.220 (0.287), 0.271 (0.371) | 0.316 (0.368), 0.372 (0.431) | 0.171 (0.274), 0.201 (0.329) |
| Average B-value ($Å^2$) | 54.2 | 89.3 | 39.1 |
| Protein | 53.9 | 89.4 | 37.7 |
| Ligand | ----- | ----- | 61.3 |
| Solvent | 62.2 | 77.1 | 53.5 |
| Overall B factor from Wilson Plot ($Å^2$) | 41.0 | 73.8 | 28.4 |
| Number of atoms | 1489 | 1185 | 1603 |
| Protein | 1440 | 1175 | 1464 |
| Ligand | 0 | 0 | 7 |
| Water | 49 | 10 | 132 |
| Protein residues | 178 | 168 | 181 |
| RMS (bonds) (Å) | 0.008 | 0.003 | 0.006 |
| RMS (angles) (°) | 0.900 | 0.560 | 0.770 |
| Ramachandran favored (%) | 97.1 | 93.6 | 100.0 |
| Ramachandran allowed (%) | 2.9 | 6.4 | 0.0 |
| Ramachandran outliers (%) | 0.0 | 0.0 | 0.0 |
| Rotamer outliers (%) | 0.0 | 0.0 | 0.0 |
| Clashscore | 8.2 | 5.3 | 3.9 |

[a] Statistics for the highest resolution shell are shown in parentheses.

Table S7. Comparison of structural features between Rosetta models and three-dimensional structures.

| Design round | Base designs | | | | | First-round designs: Internal Core | | | | First-round designs: Bottom Core | | | | First-round designs: Top Core | | | Second-round designs | | | | Third-round design |
|---|---|---|---|---|---|---|---|---|---|---|---|---|---|---|---|---|---|---|---|---|---|
| **Property / *de novo* TIM barrel** | sTIM11 | | sTIM11noCys | | DeNovoTIM0 | DeNovoTIM1 | DeNovoTIM2 | DeNovoTIM3 | DeNovoTIM4 | DeNovoTIM5 | DeNovoTIM6 | | DeNovoTIM7 | DeNovoTIM8 | DeNovoTIM9 | DeNovoTIM10 | DeNovoTIM11 | DeNovoTIM12 | DeNovoTIM13 | | DeNovoTIM14 |
| | model | structure | model | structure | model | model | model | model | model | model | model | structure | model | model | model | model | model | model | model | structure | model |
| **Structural properties** | | | | | | | | | | | | | | | | | | | | | |
| H-bonds | 236 | 204 | 194 | 171 | 198 | 201 | 197 | 201 | 201 | 209 | 176 | 138 | 189 | 201 | 184 | 177 | 169 | 174 | 167 | 195 | 163 |
| Salt bridges | 47 | 22 | 37 | 16 | 27 | 39 | 39 | 41 | 39 | 47 | 27 | 4 | 37 | 35 | 25 | 27 | 34 | 14 | 22 | 18 | 28 |
| Total area in hydrophopic clusters (Å$^2$) | 4088 | 3765 | 4260 | 4215 | 4793 | 6530 | 4800 | 4774 | 6042 | 5221 | 5195 | 5097 | 4533 | 6849 | 5812 | 5507 | 4485 | 7471 | 6381 | 6148 | 7730 |
| Area of the major hydrophopic cluster (Å$^2$) | 1171 | 1116 | 1035 | 1111 | 1066 | 2605 | 983 | 984 | 2114 | 1169 | 1180 | 1111 | 1162 | 4610 | 4611 | 4329 | 2524 | 4265 | 4416 | 4351 | 4519 |
| % of total area present in the major hydrophopic cluster | 29 | 30 | 24 | 26 | 22 | 40 | 20 | 21 | 35 | 22 | 23 | 22 | 26 | 67 | 79 | 79 | 56 | 57 | 69 | 71 | 58 |
| Total residues in hydrophobic clusters | 31 | 31 | 32 | 32 | 36 | 42 | 36 | 36 | 40 | 40 | 41 | 43 | 36 | 52 | 40 | 40 | 50 | 57 | 56 | 60 | 64 |
| Residues in the major hydrophobic cluster | 8 | 8 | 8 | 8 | 8 | 14 | 7 | 7 | 12 | 8 | 8 | 9 | 8 | 32 | 32 | 32 | 14 | 32 | 33 | 38 | 34 |
| % of total residues present in the major hydrophobic cluster | 26 | 26 | 25 | 25 | 22 | 33 | 19 | 19 | 30 | 20 | 20 | 21 | 22 | 62 | 80 | 80 | 28 | 56 | 59 | 63 | 53 |
| Total contacts in hydrophobic clusters | 82 | 81 | 89 | 91 | 98 | 120 | 84 | 88 | 112 | 104 | 103 | 113 | 94 | 143 | 122 | 114 | 129 | 152 | 133 | 150 | 166 |
| Contacts in the major hydrophobic cluster | 24 | 23 | 24 | 23 | 23 | 48 | 18 | 18 | 40 | 24 | 23 | 24 | 23 | 96 | 99 | 92 | 48 | 91 | 94 | 111 | 104 |
| % of total contacts present in the major hydrophobic cluster | 29 | 28 | 27 | 25 | 23 | 40 | 21 | 20 | 36 | 23 | 22 | 21 | 24 | 67 | 81 | 81 | 37 | 60 | 71 | 74 | 63 |
| Total ASA of the folded protein (Å$^2$) | 8661 | 8669 | 8784 | 8761 | 8920 | 8336 | 8366 | 8362 | 8360 | 8323 | 9126 | 9052 | 9149 | 8472 | 9187 | 9140 | 9002 | 9016 | 9070 | 8867 | 8730 |
| Hydrophobic ASA of the folded protein (%) | 48 | 51 | 46 | 51 | 47 | 46 | 45 | 46 | 46 | 47 | 52 | 58 | 50 | 46 | 48 | 48 | 50 | 50 | 53 | 53 | 51 |
| Total ΔASA buried on protein folding (Å$^2$) | 18893 | 17815 | 18844 | 17331 | 18199 | 19380 | 19335 | 19017 | 19429 | 19060 | 18878 | 17251 | 18141 | 18955 | 18000 | 17947 | 19469 | 19008 | 19111 | 18928 | 20048 |
| Hydrophobic ΔASA buried on protein folding (%) | 79 | 77 | 79 | 74 | 77 | 79 | 78 | 78 | 79 | 79 | 79 | 70 | 81 | 80 | 78 | 80 | 77 | 80 | 79 | 75 | 78 |
| **Physicochemical amino acid properties (%)** | | | | | | | | | | | | | | | | | | | | | |
| Aliphatic (AGILPV) | 34 | | 35 | | 37 | 38 | 35 | 35 | 37 | 39 | 41 | | 39 | 43 | 41 | 41 | 42 | 45 | 48 | | 49 |
| Aromatic (FHWY) | 9 | | 9 | | 7 | 8 | 8 | 7 | 8 | 7 | 7 | | 7 | 2 | 4 | 2 | 8 | 3 | 2 | | 3 |
| Charged (DEHKR) | 43 | | 43 | | 43 | 41 | 41 | 42 | 41 | 41 | 41 | | 46 | 39 | 39 | 41 | 39 | 37 | 37 | | 35 |
| Hydrophobic (CFILMVW) | 24 | | 24 | | 24 | 28 | 27 | 25 | 28 | 26 | 28 | | 26 | 33 | 24 | 28 | 33 | 37 | 37 | | 41 |
| Polar (DEKNQR) | 49 | | 50 | | 50 | 48 | 49 | 49 | 48 | 48 | 48 | | 50 | 46 | 48 | 48 | 46 | 43 | 43 | | 41 |
